# Robust bidirectional communication between electronics and an engineered multi-functional microbial community

**DOI:** 10.1101/2020.07.08.194043

**Authors:** Jessica L. Terrell, Tanya Tschirhart, Justin P. Jahnke, Kristina Stephens, Yi Liu, Hong Dong, Margaret M. Hurley, Maria Pozo, Ryan McKay, Chen Yu Tsao, Hsuan-Chen Wu, Gary Vora, Gregory F. Payne, Dimitra N. Stratis-Cullum, William E. Bentley

## Abstract

We developed a bidirectional bioelectronic communication system that is enabled by a redox signal transduction modality to exchange information between a living cell-embedded bioelectronics interface and an engineered microbial network. A naturally communicating three-member microbial network is “plugged into” an external electronic system that interrogates and controls biological function in real time. First, electrode-generated redox molecules are programmed to activate gene expression in an engineered population of electrode-attached bacterial cells. These cells interpret and translate electronic signals and then transmit this information biologically by producing quorum sensing molecules that are, in turn, interpreted by a planktonic co-culture. The propagated molecular communication drives expression and secretion of a therapeutic peptide from one strain and, simultaneously, enables direct electronic feedback from the second strain thus enabling real time electronic verification of biological signal propagation. Overall, we show how this multi-functional bioelectronic platform, termed BioLAN, reliably facilitates on-demand bioelectronic communication and concurrently performs programmed tasks.

## Introduction

Communication, the sending and receiving of information, drives coordinated function across a variety of systems and at many scales. Most distinct are natural biological processes and electronic technologies, with notably dissimilar communication modalities. Biological information is often transmitted as diffusion gradients of ions and small molecules. By contrast, electronic systems require electron flow, and are often wirelessly networked to extend digital communication and capabilities beyond single devices^1^. It follows that if a robust hybrid communication modality were developed, biological and electronic systems could be similarly networked in next-generation bioelectronics platforms^2^. We suggest that a redox modality can serve as this bridge.

Biological processes have been successfully explored using the ionic signaling modality to link with electronics; more recent strategies, including engineered optogenetics, innately-equipped electrogenic microbes, and/or recombinant metabolic components have enabled biological access to optical and electronic signaling^3–7^. As a more widely accessible alternative, redox-active molecules have recently been introduced as a bioelectronic signaling medium^8, 9^. Redox-active molecules are well suited for bioelectronic communication by serving dually as electron carriers and biomolecular species, where electron flow is coupled to interconversion between biologically-distinct redox states^10^. Routine electrochemical instrumentation provides a programmable electronic interface, where redox events are directed in real-time by voltage parameters and use minimal infrastructure for versatile, miniaturizable, and *in situ* implementation^11^. This provides an expedient tool to electronically access a wide repertoire of biomolecules and redox-sensitive proteins, thus potentially controlling biorecognition and associated genetic machinery^12, 13^.

Tschirhart et al. have previously realized the concept of ‘electrogenetics’^14^, where electronic input was relayed using a redox mediator to dynamically activate gene expression in *E. coli*^15^. In the present work, electrode-generated redox signals, without mediators and with surface-engineered bacteria, enable a signal-initiating transduction interface and, when coupled with an orthogonal redox mediator enabled recognition interface, bidirectional information flow between an electronic system and a community of engineered microbial cells, referred to as a BioLAN (**Fig. 1a**), is created. We use the principles of synthetic biology^16^ to coopt native redox signaling pathways for modular circuitry that coordinates information processing across populations^17, 18^.

**Fig. 1.**
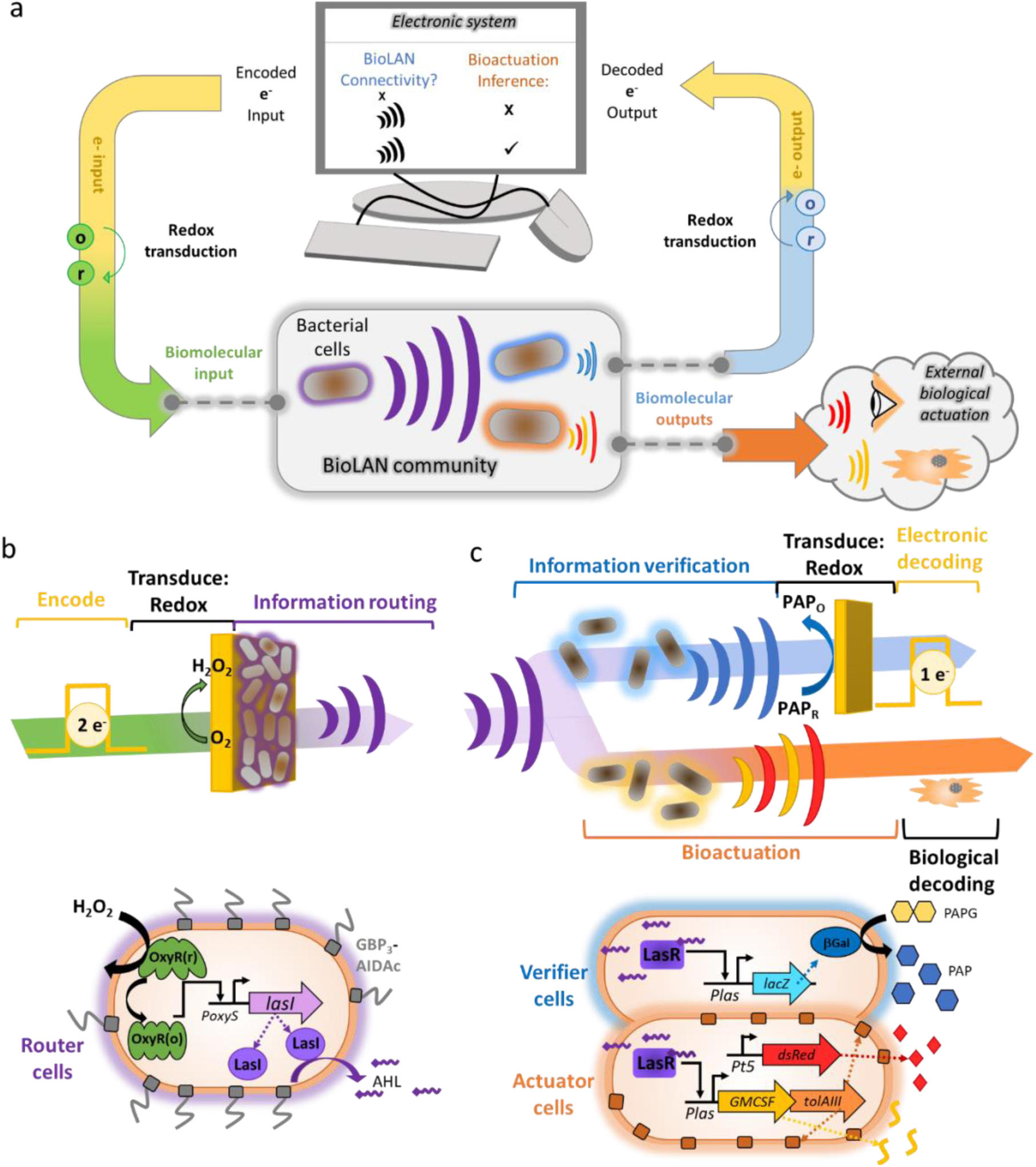
Information flow through an electronically-interfaced biological network. **(a)** Connectivity between electronics and a microbial community is established using biomolecular redox events to transduce bioelectronic signaling by simultaneously mediating electron flow and eliciting biological interactions. The result is an electronically-controlled biological local area network (BioLAN) that propagates the signal across microbial subpopulations and onto to external environments for multiplexed actuation. One subpopulation transduces received information into electronic output that returns BioLAN system status and a second produces biological outputs. **(b)** Electronic-encoded input is transduced to a biologically-recognized signal, hydrogen peroxide, via electrochemically-controlled oxygen reduction.

The electrochemically “plugged-in” BioLAN serves as an embedded, biological local area network that transmits electronic information. The BioLAN’s community nature provides several advantages – signal propagation, spatial organization, and division of labor across populations^19, 20^. Each member is engineered for specific signal processing tasks – information Routing, signal Verification, and bio-Actuation (**Fig. 1**). Initially, electronic input is redox-gated for electronic-to-bio signal transduction. The electrochemical reduction of molecular oxygen to hydrogen peroxide (henceforth, ‘peroxide’) is paired with a peroxide-inducible genetic circuit derived from the OxyR oxidative stress response regulon^21^. Once repurposed as an electrogenetic gate, cells are able to translate redox input into an orthogonal molecular cue (acyl homoserine lactone (AHL) from *Pseudomonas aeruginosa*) for information routing (**Fig. 1b**). Importantly, these Routing cells are “hardwired” to the active electrode via engineered surface adhesion. This enables direct, uniform interception of the peroxide signal and its transmission as AHL to the remaining BioLAN constituents.

The BioLAN includes two mobile populations of AHL recipients that interpret and relay signals downstream; thus a signal pathway from “encoded” electronic input to multiplexed genetic activation is established via AHL (**Fig. 1c**). Verifier cells evaluate the strength of relayed information and provide an electronically-measured response through expression of β-galactosidase (β-gal) and its cleavage of 4-aminophenyl-β-D-galactopyranoside (PAPG) to the electronically-detected p-aminophenol (PAP)^22^. In parallel, Bioactuator cells are induced by AHL to secrete a hard-to-detect therapeutic, granulocyte-macrophage colony stimulating factor (GMCSF)^23^. We show that electronic outputs from Verifier cells serve to indicate successful signal propagation as well as synthesis and secretion of the therapeutic at appropriate levels by bioActuator cells. Thus, bioLAN signaling both commands and confirms delivery of GMCSF. Collectively, bidirectional communication across a bioelectronics interface is established by the redox-specific biological processing of electrochemically-encoded and decoded information. The resulting bioelectronics platform, an electronically “plugged-in” microbial BioLAN, fulfills the fundamental requirements for inclusion of remote biological features in networked biohybrid devices.

BioLAN cells assembled at the electrode (via gold-binding peptide surface display, GBP^3^-AIDAc) intercept peroxide to activate electrogenetic expression of LasI. LasI then synthesizes acyl homoserine lactone (AHL) for signal routing. **(c)** Through AHL activation, the BioLAN indirectly connects additional cells to electrogenetically-controlled events. Verifier cells detect the routed AHL signal via LasR and produce the β-galactosidase (β-gal) enzyme. β-gal cleaves PAPG to PAP, which is detected by electrochemical oxidation. This acts to transmit the propagated signal back to the electronics and thereby reports on BioLAN system connectivity. Additionally, Actuator cells respond to the AHL signal by upregulating TolAIII-mediated membrane porosity, which co-releases a therapeutically-relevant granulocyte macrophage colony stimulating factor (GMCSF) and overexpressed DsRedExpress II fluorescent protein out of the cell via diffusion.

## Results

### 1. Bioelectronic signal transduction through OxyR-based electrogenetics

To encode and transduce electronic information to cells, we developed a new electrogenetic system using the native transcriptional activator, OxyR. *E. coli* rapidly uptakes and enzymatically degrades peroxide upon exposure, and its transient intracellular presence oxidizes OxyR (OxyR(o))^24, 25^. This can elicit a strong native regulatory response, including upregulation of *oxyS* from the *PoxyS* promoter^26, 27^. Advantageously for electrogenetic system development, peroxide can be electrochemically generated from molecular oxygen under physiological conditions^28^.

For peroxide-driven gene induction, we harnessed an OxyR-regulated genetic circuit paired with constitutive OxyR expression^29^ for peroxide-induced expression of genes of interest (*sfGFP*, *lacZ*, and *lasI*) from the *PoxyS* promoter (**Fig. 2a**). We found that *E. coli* tolerated up to 100 µM of peroxide with negligible effect on cell viability; furthermore, this concentration could be rapidly depleted by the bacteria and was expected to yield near-saturated OxyR(o) levels based on modeled kinetics (**Supplementary Fig. 1**). Thus, in agreement with previous reports, 0 – 100 µM peroxide provided a benign induction range^30^.

**Fig. 2.**
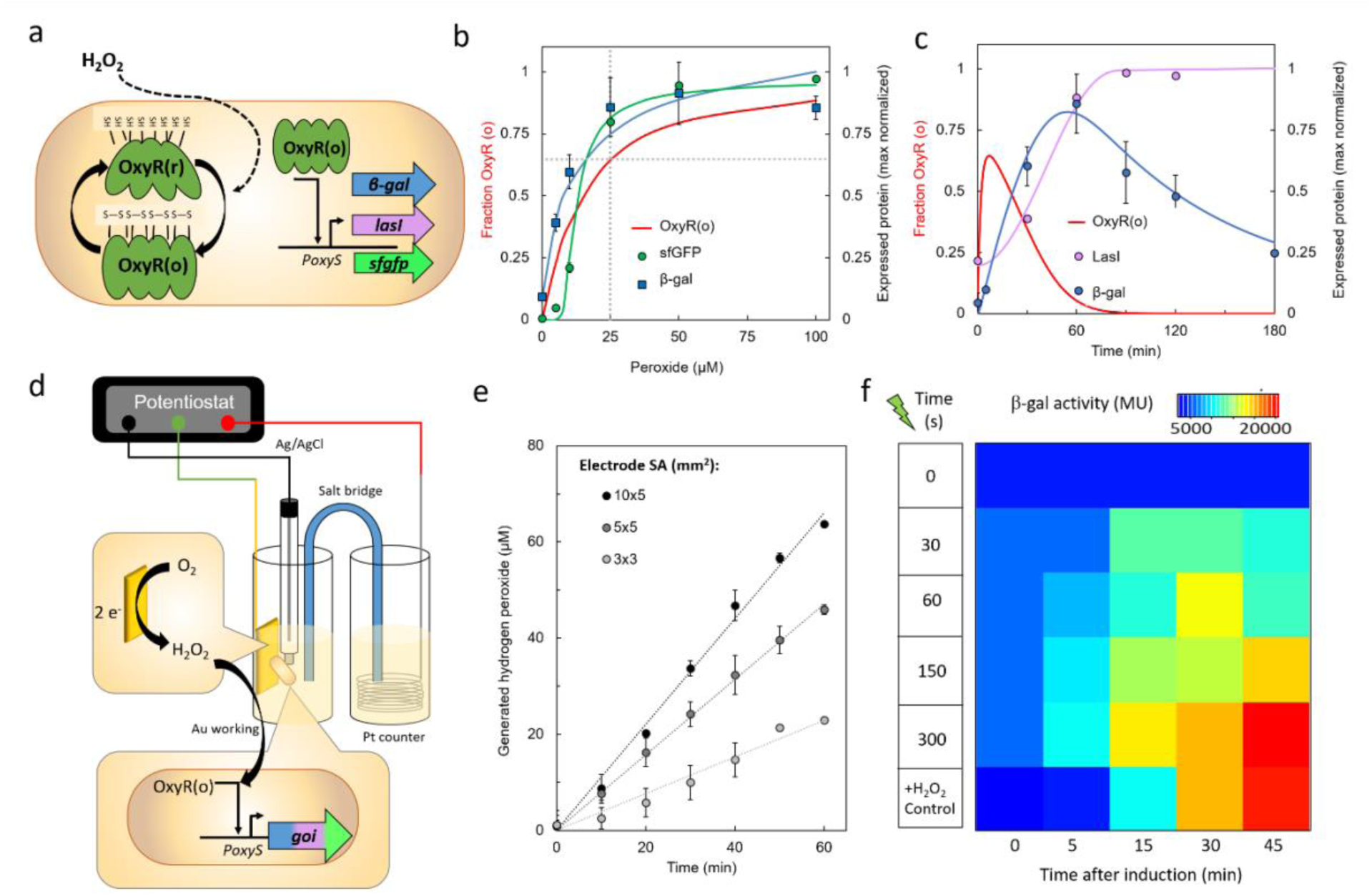
Peroxide-driven electrogenetic control. **(a)** OxyR-regulated gene expression: hydrogen peroxide oxidizes OxyR, resulting in disulfide bond formation. OxyR(o) activates expression of a gene of interest (goi) under the *PoxyS* promoter. **(b)** Dose-response of model-predicted intracellular OxyR(o) fraction at 10 min (**Supplementary** Fig. 1) and saturation-normalized expression of reporter proteins (β - galactosidase, sfGFP) to hydrogen peroxide, measured by Miller Assay (at 60 min) and flow cytometry (at 45 min), respectively. Experimental data indicated by data points. Fitted models of relative protein levels shown as solid lines. Grid lines indicate induction threshold (25 µM peroxide) and corresponding OxyR(o) peak (f_OxyR(o)_ = 0.644) required to achieve saturated protein expression. **(c)** Timecourse of model-predicted intracellular OxyR(o) fraction and data of saturation-normalized expression levels of reporter proteins (β-galactosidase, LasI, each ssrA-tagged) when induced by 25 µM peroxide, respectively. For (b-c) Experimental data are indicated by data points and fitted models of protein levels are shown as solid lines **(d)** Schematic of setup for electrochemical reduction of oxygen to hydrogen peroxide to drive OxyR(o)-mediated gene expression. **(e)** Measured peroxide concentration in the working electrode solution over time, with a constant -0.4 V applied to electrodes of indicated surface areas. **(f)** Electrogenetic β-galactosidase expression over time in response to the indicated electrochemical pulse lengths -0.4 V. ‘0 s’ condition indicates basal β-galactosidase activity. For ‘+ H_2_O_2_’ positive control dataset, 25 µM hydrogen peroxide was exogenously added to culture. Data in b and c represents means of biological triplicates and data in e represents means of technical triplicates. Error bars represent standard deviation.

To correlate OxyR oxidation state with expression levels, protein expression was modeled as a Hill function governed by a predicted OxyR(o) fraction (f^OxyR^(o)), scaled to experimental data, and correlated with f^OxyR(o)^ as a function of peroxide dose and time (**Supplementary Methods, Supplementary Figs. 2-3)**. At 25 µM, we saw near-saturated reporter expression and mostly-oxidized OxyR (peak f^OxyR(o)^ ∼ 0.6, **Fig. 2b**). Using 25 µM peroxide in a timecourse experiment, we saw LasI-generated AHL levels and β-gal peak 60 min post-induction (**Fig. 2c**), in agreement with modeled kinetics. Overall, the consistency of modeled behavior with experimental data validated the OxyR kinetic predictions. This, along with dependence of induction on cell number and protein degradation (**Supplementary Fig. 4**) provided insight toward experimental parameters.

Next we established a strategy to induce OxyR-regulated gene expression via electronic input in lieu of exogenously-added peroxide. Peroxide can be electrochemically generated at physiological conditions (pH 7) and benign voltages (approximately -0.3 to -0.9 V vs.

Ag/AgCl)^28^ by the partial reduction of oxygen, according to Equation 1.

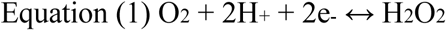

Peroxide thus provides a transduction mechanism by which electronic input can be converted into a biologically-recognized cue in our system (**Fig. 2d**). In previous electrogenetics work that coopted the SoxRS regulon, redox mediators, pyocyanin and ferricyanide, were purposely added to facilitate information exchange between the electrode and nearby cells^15, 21^. Here, in contrast, no redox mediators are added. To complete the design studies, we biased an electrode at -0.4 V for time intervals during which the detected charge transfer was stoichiometrically proportional to oxygen reduction events. Despite external influences on charge transfer (media, electrode dimensions, and oxygen availability), the correlation between charge and peroxide produced was near-constant at 58 % efficiency (**Supplementary Fig. 5**). Accumulated charge, then, presents a unit of electronic induction “dose”, with area-based proportionality between charge and peroxide production rate (**Fig. 2e**).

We tested electrogenetic induction by monitoring the β-gal activity of suspended cells, assayed periodically up to 45 min. In **Fig. 2f**, activity increases over time and as a function of charge dosage. All electro-induced samples exhibited activity above basal levels within 5 min. With a maximum dose, expression levels were on par with a 25 µM exogenously-induced control, which is indicative of having reached expression-saturating OxyR(o) levels. We saw similar charge-based induction of sfGFP levels and verified that growth effects due to electrochemically-produced peroxide did not differ from exogenously-supplied peroxide (**Supplementary Fig. 6**). Thus, this novel electrogenetic circuit enables us to link electronic input, through redox transduction and without need for mediator addition, to biological behavior.

### 2. Elucidation of the bioelectronic interface

Electrode-generated peroxide in a quiescent fluid creates a far more conducive signaling environment compared to the exogenous addition to a planktonic culture, where peroxide must be uniformly mixed. Thus, we sought to elucidate the spatiotemporal dynamics of the process to understand its effects and to find optimal conditions for uniform electrogenetic induction.

First, we simulated the peroxide gradient at the electrode in the presence of unstirred planktonic cells based on its generation rate, diffusion, and consumption by the cells (**Supplementary Methods**). The model data in **Fig. 3a** shows that peroxide reaches micromolar concentrations proximal to an electrode surface within seconds of a voltage bias through diffusion. Despite the strong peroxide consumption behavior of the cells (**Supplementary Fig. 1**), their presence as a planktonic population only marginally attenuates the bulk concentration (**Supplementary Fig. 7**). Conversely, in a scenario where the entire cell population is distributed at the electrode surface, nearly all generated peroxide is intercepted, thereby limiting its accumulation in solution to 1.5 µM at peak concentration (**Fig. 3b**). Once peroxide generation ceases, the simulation shows an interfacial zone of complete elimination by the bacterial monolayer within 1 min.

**Fig. 3.**
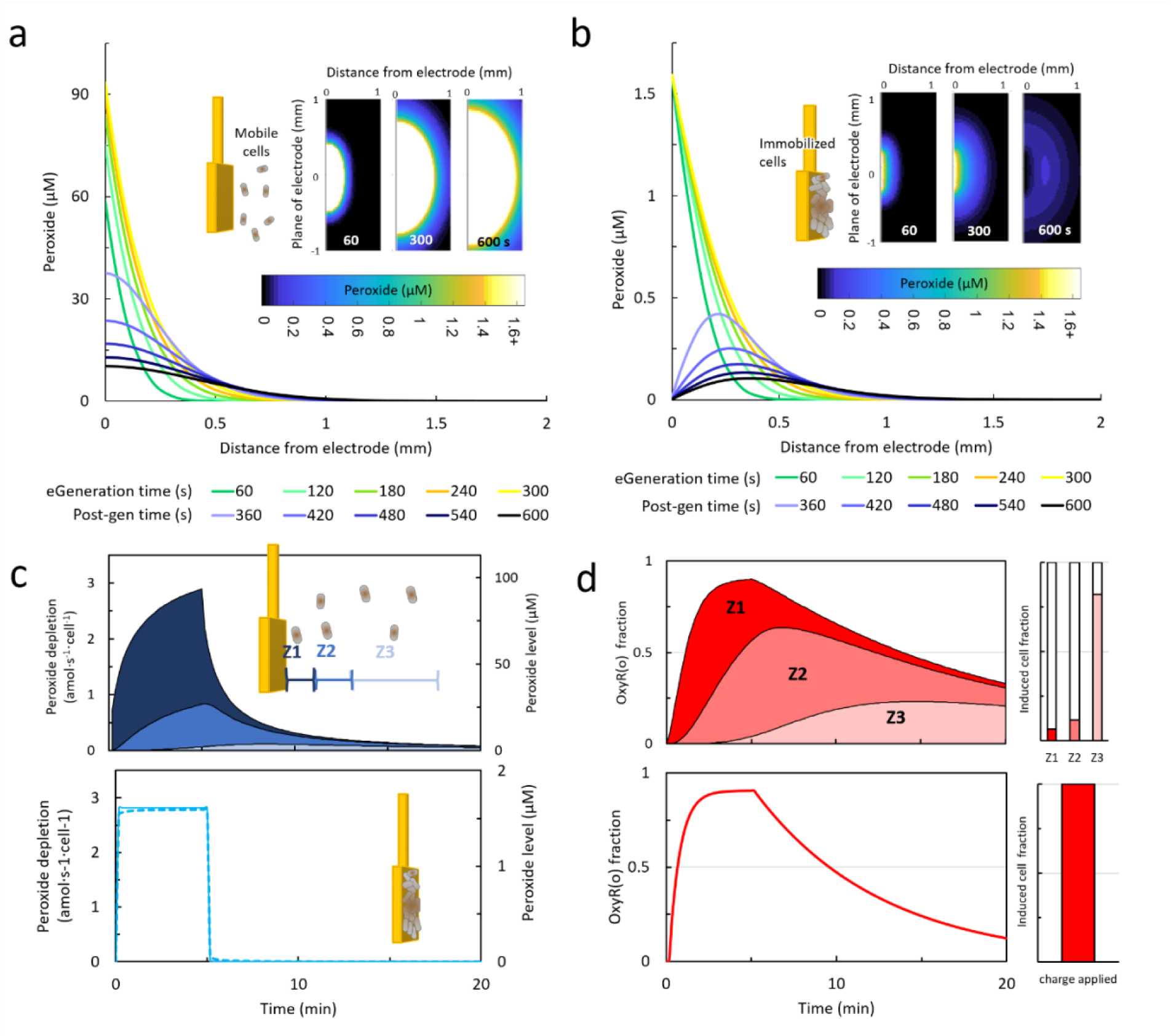
Contribution of electrochemical generation and bacterial consumption rates to hydrogen peroxide flux simulated at an electrode. **(a)** Simulation of peroxide flux at a distance from the electrode interface over time, based on surface-area dependent rate of charge, average stoichiometric efficiency of oxygen reduction, and the presence of mobile peroxide-consuming cells in the electrode’s vicinity. 0-300 s timepoints account for a biased electrode with continuous peroxide generation (“eGeneration”). For 360 – 600 s timepoints, the electrochemical reaction has been terminated. **(b)** Simulation of peroxide flux at an electrode interface, differing from (a) due to the positioning of the cells at the electrode interface. **(c)** Modeled ranges of peroxide depletion over time, predominantly by cell decomposition, based on peroxide electro-generation in the presence of mobile cells (tracked across distance intervals from electrode, Z1, Z2, and Z3), and at the electrode interface for immobilized cells. **(d)** Projected ranges of OxyR oxidation state over time for cells corresponding to the conditions denoted in **(c)**. Oxidized (OxyR(o)) levels reported as fractions of total intracellular OxyR. Additionally, the population fraction of cells applicable to the OxyR models is reported, based on cell. For planktonic cells, the peroxide depletion rates in (c) and OxyR(o) fractions in (d) are denoted as a range of values within three zones proximal to the electrode interface: Z1 (0-0.225 mm), Z2 (0.225-0.625 mm), and Z3 (0.625 – 3.5 mm).

We next analyzed the effect of the cells’ position on OxyR dynamics. In **Fig. 3c-d**, we found that the consumption by cells near the electrode greatly limits the peroxide quantity available for distal cells, whereas all electrode-localized cells provide uniform consumption. For planktonic populations in the simulation, less than 7 % of cells experience maximal OxyR activation in a peroxide-rich environment (*i.e.* at least 25 µM peroxide whereby f^OxyR(o)^ ≥ 0.6), which exists within 225 µm of the electrode surface. The peroxide concentration declines with distance, resulting in the majority of cells (82 %), which are positioned at least 625 µm away from the electrode, to experience negligible peroxide levels and minimal OxyR activation (*i.e.* less than 4 µM for f^OxyR(o)^ ≤ 0.1). By contrast, the fractions of activated OxyR and, consequently, of induced cells are higher for electrode-localized cells - achieving 0.9 f^OxyR(o)^ universally within 5 min of voltage bias (**Fig. 3d**).

Additionally, we note that these electro-induction parameters (*i.e.* immobilized configuration of the bioelectronics interface, 300 s voltage duration) yield net peroxide uptake nearly-equivalent to 25 µM induction (**Supplementary Fig. 8**), which was shown to sufficiently saturate protein expression (**Fig. 2b**). Thus, these design studies for *in situ* peroxide generation elucidate favorable conditions that should enable strong gene expression (*e.g.* for LasI-mediated AHL signaling) and, overall, provide a fast, efficient, and uniform bioelectronic information-transfer interface.

### 3. Electrode-immobilization of electrogenetic cells

In order to localize signal transfer at the bioelectronic interface, cells were engineered specifically to enable their assembly directly onto the electrode via peptide-mediated affinity interactions. We harnessed the outer membrane autotransporter pore-forming protein (AIDAc) as a vector for cell surface modification^18, 31, 32^. We fused AIDAc with a recombinant peptide consisting of a trimeric repeat of a non-natural peptide characterized for its high affinity to gold (GBP^3^)^33–36^. Our structural prediction of the GBP^3^-AIDAc fusion (**Fig. 4a**) depicts the peptide’s extrusion through the center of the AIDAc barrel - a relatively unstructured conformation that extends outward while remaining tethered at its C-terminus.

**Fig. 4.**
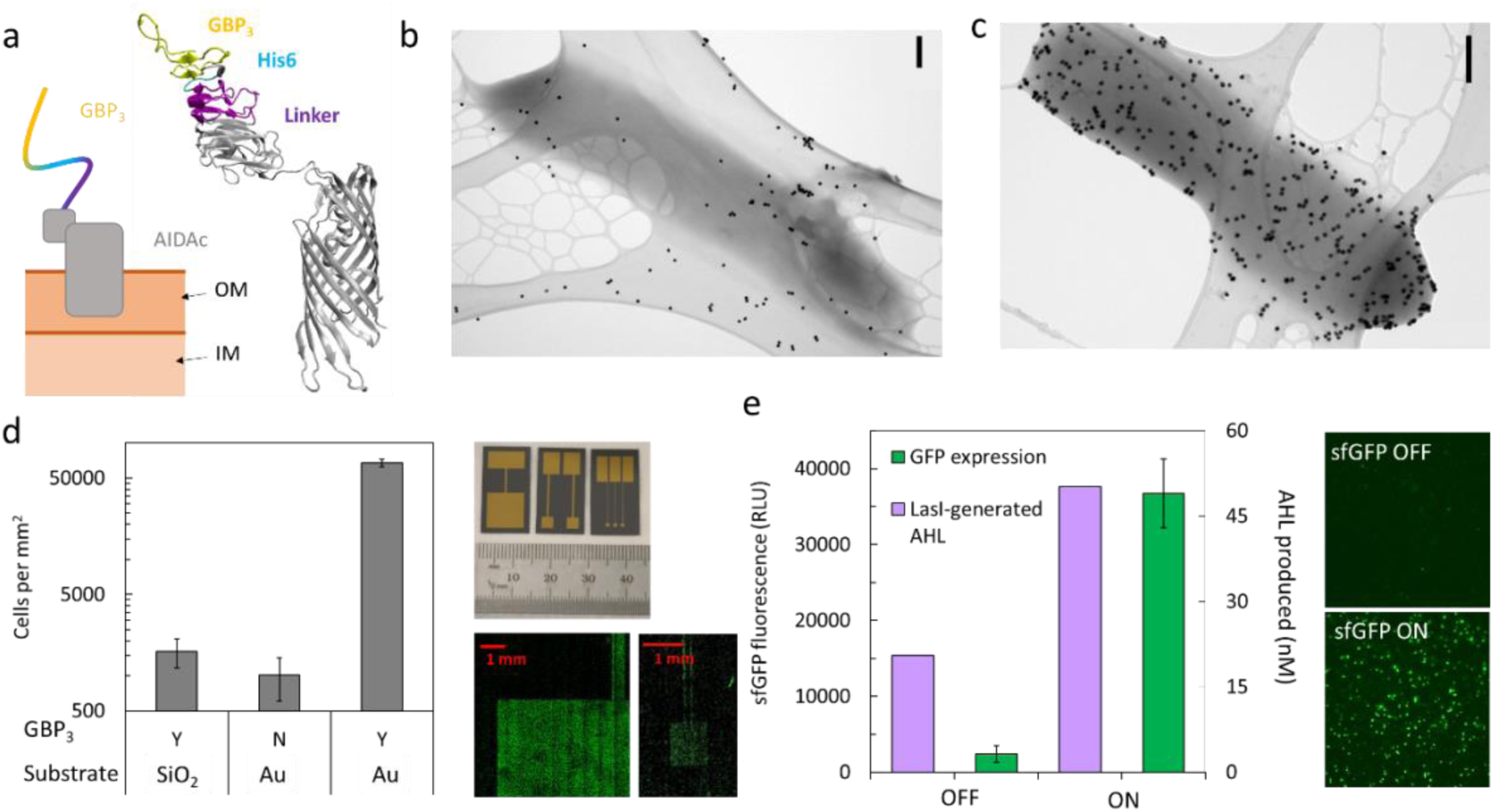
Assembly of electrogenetic cells onto gold surfaces via peptide surface display. **(a)** Homology model of AIDA autotransporter structure with an N-terminally-linked fusion of ‘GBP^3^’ gold-binding peptide and hexameric histidine tag. Transmission electron micrographs of *E. coli* cell lacking labeled with 20 nm gold nanoparticles, lacking genes for GBP^3^ surface display in **(b)**, or GBP^3+^ in **(c)**. Scale bars = 200 nm. **(d)** Coverage of *E. coli* cells (- / + pGBP^3^) immobilized to either a gold-coated (Au) or uncoated (SiO^2^) silicon wafer and stained with Syto-9. **(e)** Demonstration of electrogenetic activation of gene expression by applying a voltage to electrodes with immobilized cells hosting various reporters (AHL quantified by Curve 2 in **Supplemental** Figure 2**-2e**). Quantification for d and e are based on three biological replicates per condition and image analysis of technical replicates with standard deviation of data reported.

For initial characterization, surface-displayed expression of GBP^3^ was tested by quantum dot (QD) labeling, which showed affinity for the GBP^3+^ and not the GBP3^-^ cell surface, presumably due to the Zn-containing QD shell and the peptide’s histidine tag through well-known affinity interactions with transition metals (**Supplementary Fig. 9**)^37, 38^. Furthermore, by transmission electron microscopy, gold nanoparticles did not associate with cells lacking GBP^3^ (**Fig. 4b**), but showed retention on GBP^3+^ cell surfaces (**Fig. 4c**). We found that GBP^3+^ cells bound with fifty-fold higher specificity to planar-deposited gold compared to a silicon wafer’s native oxide. This promoted cell templating on patterned gold electrodes (**Fig. 4d**). We show representative composite fluorescence images of Syto9-stained cells immobilized on patterned gold electrodes with surface areas between 1 and 100 mm^2^ (**Fig. 4d**). Cell distribution was visibly uniform and spatially confined to reflect the geometry of the electrode. In this way, electronic information transfer to cells is uniform, predictable, and based on the electrode surface area.

Next, we confirmed that gold-bound cells could be electronically-induced to express sfGFP and LasI (through *in situ* electrochemical peroxide generation) (**Fig. 4e**) with maintained viability and statistically-similar peroxide consumption rates to those of planktonic cells (**Supplementary Fig. 10**). Overall, the surface-assembled GBP peptide establishes the physical connectivity that is important for efficient information flow across the bio-electronic interface.

### 4. Reflexive bioelectronic verification of information relayed throughout a cell network

We next aimed to propagate the electrogenetic cue across multiple cell populations by incorporating cell-to-cell communication and, moreover, to enable electrochemical verification of signal transmission by producing a redox-active output upon signal exchange. We accomplished this by networking two key cell populations, each designated for signal routing and verification, respectively (**Fig. 5a**). First, an electrogenetic system designed to produce LasI-synthesized AHL using electrode-bound cells allows for AHL signal propagation (*i.e.* signal routing) from the bioelectronic interface through diffusion. The diffusion space of AHL establishes network boundaries to which AHL-recipients can be connected, for instance, to verify signal strength. Further, as a planktonic population, these verifiers could intercept the AHL signal throughout the network space. Here, we establish such signal Verifiers to respond with electrochemical feedback, where signal strength is reflected by the cells’ genetic response and resulting electrochemical activity (**Supplementary Fig. 11**). Specifically, AHL-induced β-gal catalyzes production of electrochemically-active p-aminophenol (PAP), which is transduced into an electronic output by its electrochemical oxidation^39, 40^. Provided that the electronic output reflects the input status, continuity in information transfer across the bioelectronic interface is thus maintained bidirectionally.

**Fig 5.**
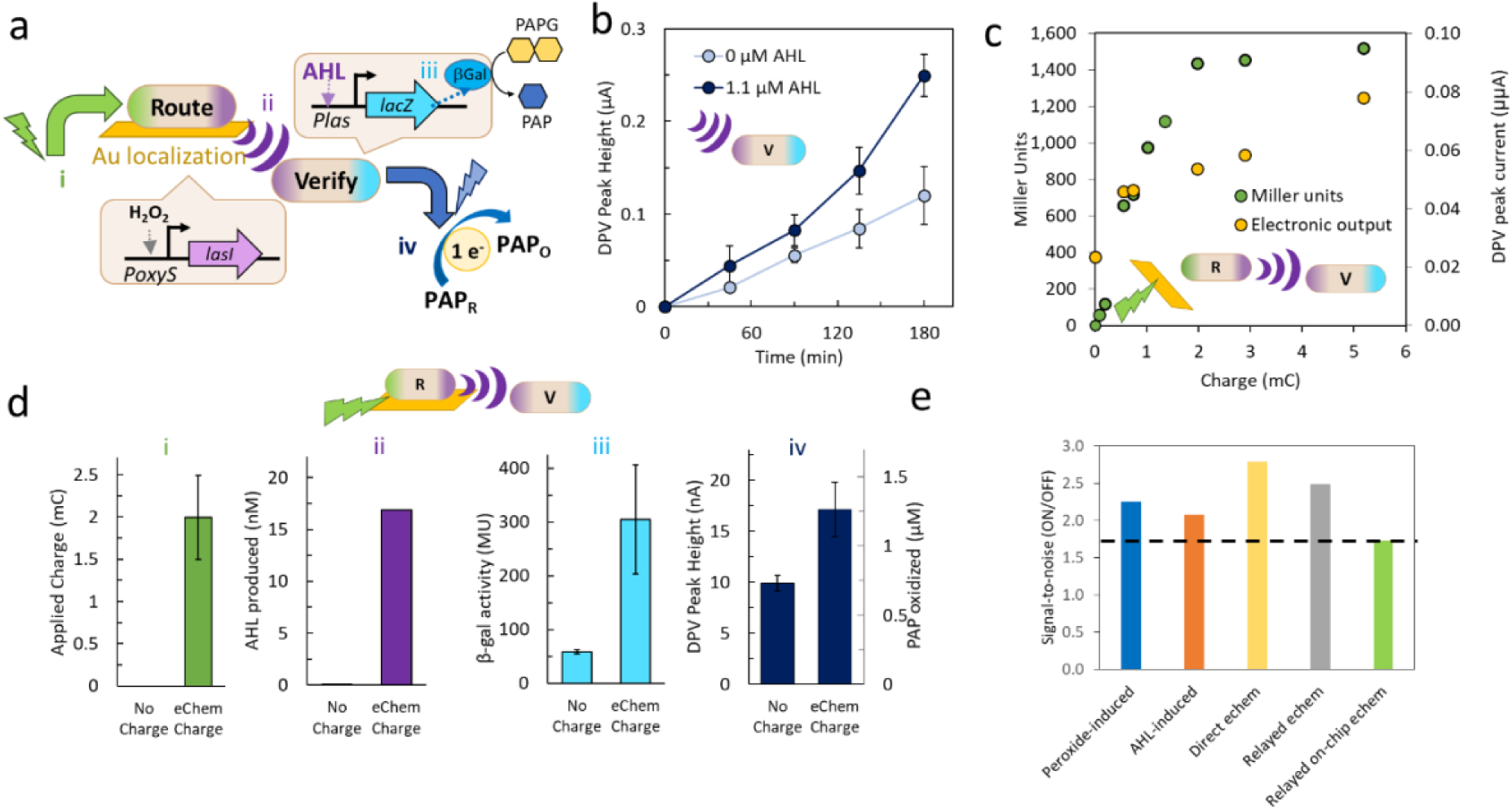
Electronic information flow through an engineered two-member community. **(a)** Schematic of information flow from the electrode, through the Routing and Verification cells with AHL, and back out to the electrode. Redox signals provide transduction between cells and electrodes. **(b)** Electronic output, quantified by DPV peak current of PAP oxidation, from Verifier cells with or without AHL induction, measured over time. **(c)** Verifier cell output, measured both electronically (PAP oxidation current) and with the Miller Assay in response to induction with different charges. Routing cells are planktonic. **(d)** Indicators of the applied charge (i): AHL production levels (ii); -gal activity (iii); and electronic output of a two-population bioelectronic system described in (a) either, with or without applied charge. **(e)** Signal-to-noise ratios (On/Off) at either 1.5 or 3 hours post-induction. Data are from Supplemental Figure 5-1 and Figure 5 b and d. In d, means of biological triplicates are shown; error bars indicate standard deviation.

We first confirmed that planktonic AHL-sensing cells could produce an electrochemically-detectable output based on β-gal cleavage of PAPG into PAP. Differential pulse voltammetry (DPV) of AHL-induced cultures in a PAP-oxidizing voltage range (0.25 to -0.15 V) revealed a faster increase in peak current over time compared to uninduced samples (**Fig. 5b**). We next included electrogenetic AHL-producers and provided charge to this co-culture. We found sustained and concomitant production of β-gal (measured via Miller assay) and PAP (measured electrochemically), both commensurate with charge “dose” (**Fig. 5c**). While the Miller Assay is performed at the time of measurement, PAPG is added to the cells immediately after electro-induction and PAP accumulates over time as the cells are cultured, allowing for continuous measurement without sample loss.

Having confirmed that electrogenetic input could be relayed from a router population to recipients, which validate charge input via electrochemical feedback, we assembled the electrogenetic AHL-producing cells onto the electrode to improve signal efficiency (as per the previous section). **Figure 5d** portrays data from the bioelectronic system with immobilized Routers and mobile Verifiers under ON and OFF conditions, where a bias was applied to activate electrogenetic response only in the ON condition (i). Electrode sizes and representative electrode coverage by Router cells were similar between the two conditions (**Supplementary Fig. 12**). Production of the intermediate signal, AHL, increased fifteen-fold in the ON condition (iii), attributed to Router cell electrogenetic activation of LasI. Additionally, the AHL-induced β-gal expression in Verifier cells was measured to have five-fold enhanced enzymatic activity in ON samples (iv). Finally, the electronic output of ON cultures was approximately two-fold higher than the OFF sample output (v).

We define the signal-to-noise ratio (SNR) as the ratio of peak DPV output signals from the ON and OFF samples. The SNR **(Fig. 5e**), in terms of electronic output, is approximately two-fold and is similar between the relayed on-chip, relayed planktonic, and a peroxide-induced alternative single-cell-type system (**Supplementary Fig. 13**). The robustness of the system is demonstrated by noting that the SNR was consistently near 2 irrespective of the means for actuating the cells (electronic or chemical induction).

Thus, we have confirmed robust bioelectronic information transfer through a bacterial community. The electronic input was initiated at an electrode, where peroxide is generated. A surface-attached electrogenetic cell layer, serving as a Router population, transduces this signal via genetic response – to a molecular signal, AHL. A suspension of planktonic responding cells (Verifiers) de-codes the AHL and synthesizes β-gal, which in turn is measured electronically via conversion of PAPG to PAP. The output is also verified using a standard Miller assay. The net result demonstrates a complete electronic-to-bio-to-electronic, and thus, bidirectional information flow.

### 5. Coupling bioelectronic information exchange to bioactuation

By delocalizing electrogenetics from a discrete surface to a multi-population community through AHL communication, the bioelectronic system exploits native biological signaling processes to accommodate non-electrogenetic cell types, all while electronic surveillance is maintained (from PAP electrochemistry) to verify cross-population information transfer. It follows then that numerous and distinct populations could be connected in: for example, the inclusion of AHL-responsive cells designed to perform biological functions allows for task-execution in parallel with electronic feedback from Verifier cells. Thus, we designed an AHL-inducible Actuator response, where designated cells perform an executive function when turned ON. As the intended function, Actuator cells express and secrete protein products that, in turn, might influence environments outside of the AHL-networked bioelectronics. Specifically, AHL-inducible cells co-secrete a natively-recognized biotherapeutic peptide, granulocyte macrophage colony-stimulating factor (GMCSF) and DsRed as a fluorescent marker (**Fig. 6a, Supplementary Fig. 14**). GMCSF has therapeutic efficacy for Crohn’s disease; its secretion by *E. coli* has been previously accomplished at physiologically-relevant levels by co-expression of the TolAIII pore.^23^ In conjunction, DsRed provides optical reporting of secretion.

**Fig 6.**
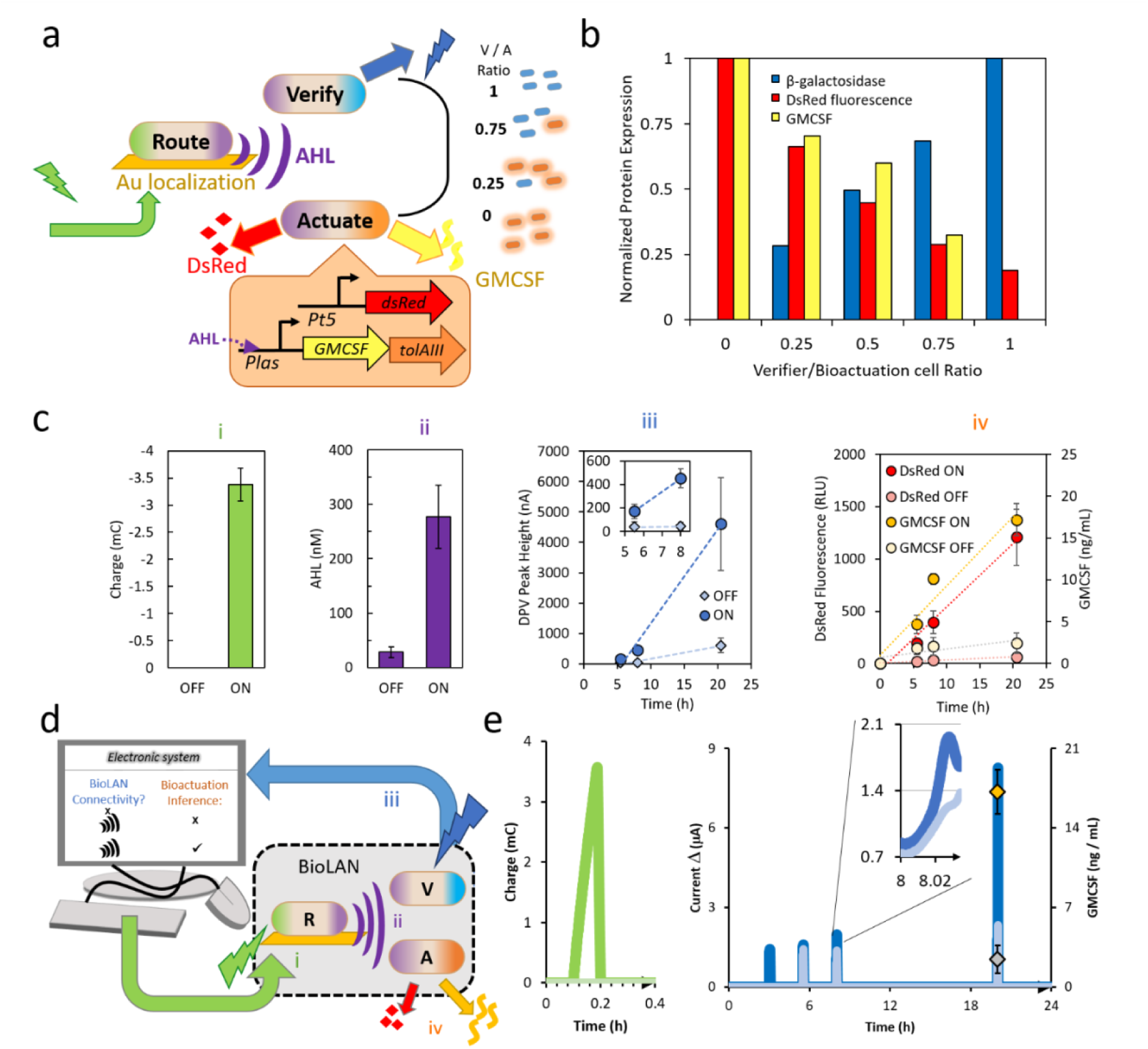
BioLAN function. (a) Schematic of information flow into the BioLAN, communication between members, and BioLAN output. (b) Normalized protein expression from BioLAN communities in which the indicated ratios of Verifier/Actuator cell ratios were used. (c) Measurements of applied charge dose (i), AHL production levels (ii), electronic output from Verifier cells (iii) and GMCSF and DsRed production by Actuator cells (iV) from BioLAN systems induced electronically (ON) or kept uninduced (OFF). Means and standard deviations of biological quadruplicates are shown. (d) Schematic showing BioLAN connection to a computer and information flow between them. (e) Representative traces of real-time measurements of input charge (i) and output current (ii), which precedes the detection of Actuator cell production of DsRed and GMCSF (diamonds). Data is representative and corresponds to that in (c).

We created consortia of varied composition of Router, Verifier, and Actuator cells and probed for their diverse outputs (β-gal, extracellular DsRed and GMCSF) upon AHL signaling. We found that the AHL-driven outputs strongly correlated with the respective population ratios (**Fig. 6b**), which demonstrates the diversity and resolution of networked functions. Hence, in addition to electrode size, applied charge, and genetic circuits introduced to the Router cells, the population ratio could be used as yet another tunable parameter for multiplexed consortium function.

To illustrate the fully-networked bioelectronics system, we established electrochemical connectivity by pairing electrode-bound Router cells with the co-culture of planktonic Verifier and Actuator cells. Samples were either maintained at an OFF state or turned ON with about -3.5 mC (**Fig. 6c (i)**). To track the activity of the community, we measured AHL levels (**ii**), electrochemical output (**iii**), and secretion levels (DsRed and GMCSF) (**iv**) throughout the incubation period. All of the outputs show similar trends for OFF vs ON samples. In OFF samples, the AHL, PAP, DsRed, and GMCSF levels remained low throughout a 20 hour period. Conversely, in electronically-induced communities, levels of measured outputs increased dramatically and show 4 to 10-fold higher levels than those in OFF samples. Interestingly, in this configuration, all outputs showed a difference between the ON and OFF conditions after 5.5 hours and continued to diverge over time. This demonstrates for the first time bioelectronic I/O successfully directing and reporting distributed biological task execution in a three member microbial community through an electrochemically-triggered and -read molecular information relay.

This particular consortium grouping is uniquely suited to function as a biological local area network (BioLAN) by bioelectronic information exchange between the electronic I/O and AHL-delineated coordination across Router, Verifier, and Actuator cells (**Fig. 6d**). That is, cell-to-cell communication as exemplified in this BioLAN, enables distribution of functions into a collective that is modular and multiplexed by cell type. Further, the consortium members developed here expand the communication repertoire by including electrode-synthesized redox-transducing electronic signals (peroxide, PAP), resulting in convenient interfacing capability with an electronics infrastructure. The electronic information flow can clearly be traced through induction current (**Fig. 6e (i)**) to the BioLAN output current **(ii)** to confirm system connectivity as real-time electronic feedback. This, in turn, informs on therapeutic secretion status due to AHL-correlated responses (**Fig. 6e (ii)**).

## Discussion

This work enables bioelectronic interfacing with a living biological network (BioLAN) by purposing redox molecules to interconvert between electronic and biological I/O to achieve bidirectional information exchange. Redox signal transduction provides “wiring” to connect to the synthetic biology, whose electronic proficiency is established by minimal genetic rewiring. The engineered cells of the BioLAN have genetically distinct roles of signal routing, verification, and actuation and remain interconnected via internal AHL communication. The BioLAN “plugs into” the electronics infrastructure to seamlessly transmit information from electronics, through biology, and back out to electronics, and to concurrently drive programmed biological function.

The BioLAN processes electronically-encoded information using redox. As hydrogen peroxide is a universally-recognized redox molecule in biology that can be electrochemically-generated, we repurposed native oxidative stress regulation for electrogenetic expression via the OxyR protein and PoxyS promoter. In contrast to the previously-developed SoxR-based electrogenetic circuit, the newly-characterized system functions in aerobic environments and eliminates the need for addition of external mediators. Correlation between modeled peroxide-oxidized OxyR and charge input ensures that electrochemically-supplied peroxide is targeted at non-toxic, yet above-threshold levels for gene expression. Ultimately, voltage-mediated control over the electrogenetic response encodes an ON or OFF status onto the BioLAN. The electrogenetic cells serve as a Router component to distribute the information from the electronics throughout the BioLAN. This is accomplished by system-specific signal propagation from the electrogenetic cells, which transduce the peroxide input signal into an orthogonal output, AHL, via induction of its synthase. It is important to specifically note the AHL communication network is orthogonal to the microbes involved – they do not respond independently to AHLs. Should native signaling processes be desired, the synthesis (and secretion) of other molecules could be engineered into the Router cells. Bacterial autoinducer AI-2, for example, would enable natural processes to bias the remaining cells in the BioLAN^41^.

Also, electrode localization of Router cells improves *in situ* induction kinetics and avoids diffusion limitations that can lead to heterogeneous responses in suspended biological systems, as confirmed by modeling OxyR redox state^42^. Together, we show that the generation of an electrode-localized peroxide flux, its use for bio-electrochemical information transfer, and its pairing with the surface-attached cells, yields a robust, quasi-solid state electrogenetic platform. This setup invites future opportunities for scalable, spatially-programmable biological control via three-dimensional, soft, flexible, miniaturized and arrayed electrode formats ^43–47^. By combining this “hardwired” format with “wireless” signal propagation, one can envision AHL providing a high fidelity Routing capability that extends the network’s boundaries to remotely-located AHL-responders Given that wireless network components typically return signals to confirm connectivity, the BioLAN should include reflexive feedback. That is, connectivity should prompt AHL-responders to provide a verification signal directed to the central electronics. Here, Verifier cells inform on AHL-connectivity via PAP redox output; being mobile, they reflexively signal across the physical boundaries of the network with output signal amplitude as a real-time indicator of connectivity strength. In demonstrations, electronic feedback was measurable for at least 20 hours post-induction, and provided an ON/OFF system verification within 3 hours.

Finally, the peripheral BioLAN components should perform electronically-programmed functions and thus enable remote bioelectronic actuation. Importantly, because Actuator cells homogenously co-populate the BioLAN with the Verifiers and utilize the same AHL-sensitive circuitry for gene expression, the Verifiers’ redox-activity infers the status of co-occurring actuation events. Here, release of DsRed and GMCSF payloads accumulated proportionally to electrochemical readout over time, thereby validating the electronic output as a proxy of secretion status. Furthermore, actuation offers extended connectivity to external environments based on signal recognition: extracellular fluorescence (DsRed) as optical readout and a secreted biologic for potential *in vivo* interactions (GMCSF). Signal fidelity across all transduction formats was maintained and was tunable based on population ratio. In this way, the networking of multiple AHL-responsive cell types expands the influence of electrogenetic control, multiplexes outputs, and enables the electronic orchestration of complex tasks through signal relay, spatial segregation, and division of labor.

Overall, because redox is both accessed electrochemically and serves as a medium of information exchange throughout biology, this communication mode presents a reliable platform for “plugging in” the bioLAN to electronic devices. The established bidirectional dialogue yields a hybrid system with electronically-programmed and tracked, yet biologically-executed, functions. Bioelectronics connectivity featuring synthetic biology may contribute to a future Internet-of-BioNano-Things^48^, yielding applications capable of embedding “biological intelligence” in ecological settings, wearable interfaces, and *in vivo* environments.

## Methods

### Chemicals

4-Aminophenyl β-d-galactopyranoside (PAPG), 4-aminophenol (PAP), and ortho-Nitrophenyl-β-galactoside (ONPG), propidium iodide (PI) were from Sigma-Aldrich. PAPG was dissolved in diH2O and PAP and ONPG were dissolved in 0.1 M phosphate buffer (PB). Agar, KCl, LB broth (Miller), M9 salts, glucose, glycerol, casamino acids, magnesium sulfate (MgSO_4_), calcium chloride (CaCl_2_), 3-(N-morpholino)propane sulfonic acid (MOPS), and hydrogen peroxide (H_2_O_2_, 30%) were from Sigma Aldrich. N-3-oxo-dodecanoyl-L-Homoserine lactone (AHL) from Cayman Chemicals. Various biological fixatives included formaldehyde (ThermoFisher Scientific) and paraformaldehyde (Electron Microscopy Sciences).

### Cell culture and media

Unless otherwise indicated, cells were grown overnight in LB at 37 °C aerobically with 250 rpm shaking, and then inoculated from the overnight cultures at 1-2% in LB or M9 media and grown until the indicated cell density (OD600). M9 media consisted of 1 × M9 salts, 0.4% glucose, 0.2% casamino acids, 2 mM MgSO_4_, 0.1 mM CaCl_2_ and, optionally, 100 mM MOPS.

### Molecular biology and genetic engineering

All enzymes, competent cells, and reagents were purchased from New England Biolabs (NEB) and used according to provided protocols. Polymerase chain reactions (PCR) for DNA amplification used Q5 polymerase and oligomeric primer sequences denoted in **Supplemental Table 4**. Post-PCR DpnI digestion of plasmid templates, polynucleotide kinase phosphorylation, T4 ligations, and Gibson fragment assembly, and chemical transformation into *E. coli* were performed using NEB products and standard molecular biology techniques. DNA clean-up, gel extraction, and plasmid prep kits were purchased from Zymo Research and used according to provided protocols. Synthetic gene fragments GBP3 and proD were purchased from Integrated DNA Technologies (IDT). All final plasmid constructs were sequence-verified. Constructs included pOxyRS-sfGFP-laa, pOxyRS-LacZ-laa, pOxyRS-LacZ, pOxyRS-LasI-laa, pOxyRS-LasI, pBla-GBP3, pBla-His6-AIDA, pBla-Linker-AIDA, pAHL-LacZ, and pAHL-GMCSF.

### Binding of GBP^3+^ cells on chip

Cells were grown overnight in LB medium, then diluted in fresh medium (as indicated) and grown until mid-log OD (unless otherwise indicated). Electrodes or pre-cut gold chips were cleaned by transferring from acetone to isopropanol to water solutions. These were then dried with a nitrogen gas stream and subjected to ozone treatment for 1 h using a PSD Series UV ozone system (Novascan Technologies, Boone, IA). The immobilization protocol was customized to each strain to reliably yield at least 40 % surface coverage. Small volumes (50-100 µL) of cultures were either applied directly to the surfaces of face-up gold chips to achieve edge-to-edge surface coverage, or alternatively, first centrifuged (8000 rpm, 3 min) to concentrate approximately 4-fold prior to application. After a 30 minute RT static incubation, the cells were washed in PBS face up (each chip placed inside a well of a 6-well culture plate) for 15-30 minutes at 150 rpm. Cell-covered electrode chips were incubated in PBS at RT until further analysis or induction.

### Fluorescent cell staining

To visualize electrode-immobilized bacteria, cells were fluorescently stained using a Live/Dead Baclight Bacterial Viability Kit (Molecular Probes) in preparation for fluorescence microscopy imaging. The viability kit’s dyes, Syto9 and propidium iodide (PI), were used alone or in combination for various experiments. In each case, the reagents were diluted between 667 and 1000-fold in PBS, then applied to the surfaces of cell-covered gold chips and incubated in the dark for 45 min, undisturbed, to stain immobilized cells, after which the chips were rinsed and stored in PBS, prior to fluorescence microscopy and image analysis, as described below. Initial characterization of surface display protein sequence (Linker-AIDA, His-AIDA, or GBP3-AIDA protein) contribution to bacterial immobilization was determined by live cell imaging, where only the Syto9 stain was applied. For subsequent analyses of immobilized cell density, a fixative solution (prepared at 2 %) was first applied for 30 min to preserve the samples, followed by a PBS rinse and application of a PI solution prior to imaging. To quantify viability of cells bound to electrode chips, Syto9 and PI were applied in combination, followed by the fixative solution, with PBS rinsing in between steps. In this case, both green and red fluorescence images were captured as described below. To calculate percent of dead cells, the total cell area of PI-stained cells in the image was divided by the PI + Syto-9 stained cell area. The fluorescence images were converted to 8-bit, consistently thresholded, and the cell areas were calculated using ImageJ (NIH). For dead-only staining of planktonic cultures, PI was used at 0.5 mg/ml in 0.85% NaCl. After 0.5 - 2 mL of the cell culture (depending on cell density) was pelleted, 50 µl of the PI solution was added, the cells were re-suspended, and incubated in the dark for 30 minutes. After a PBS rinse, cell fluorescence was read on the flow cytometer or imaged by the fluorescence microscope as described below.

### Fluorescence microscopy

An Olympus BX53 microscope with the 49002-BX3; ET-EGFP/FITC/CY2 470/40X, BS495, 525/50M filter cube was used to visualize sfGFP-producing and cells stained with Syto9 fluorescent dye. For propidium iodide-stained cells the 49008-BX3; ET-MCHERRY/TXRED 560/40X BS585, 630/75M filter cube was used. Additional imaging utilized either an Olumpus BX60 for green fluorescence or a Zeiss Axio Observer 7 (Carl Zeiss AG, Oberkochen, Germany) with Colibri LED illumination and a *Filter Set* 43 HE *Cy3* for red fluorescence. Composite fluorescence images were obtained using a Zeiss LSM700 confocal microscope.

### Peroxide electrochemical setup and generation

For electrochemical peroxide generation a gold-patterned silicone wafer chip (with electrode dimensions 1 cm^2^, unless otherwise indicated) was used as a working electrode. A coiled platinum wire (BaSi) with surface area larger than that of the working electrode was used as the counter electrode. An Ag/AgCl (BaSi) reference electrode was used. Using sufficient volumes of the indicated solution or media, the electrodes were completely submerged, with the working and reference electrodes in a separate glass vial than the counter electrode, but connected by two salt bridges. The working electrode solution was undisturbed (or, where indicated for comparison, stirred with a 7mm stir bar and a mini magnetic stirrer). The electrodes were connected to a potentiostat (either 700-series CH Instruments or BioLogic VMP3). Chronoamperometry, poised at -0.4V for the indicated duration, was performed to generate hydrogen peroxide. The endpoint charge was recorded for each run.

Agar salt bridges consisted of 6 inch-long 1.2 mm OD, 0.9 mm ID glass capillary tubes bent into a U shape after brief heating under a Bunsen burner. A 3% agar solution with 1 M potassium chloride (KCl) was heated and added into the bent capillary tube. Tubes were cooled by immersion in a 3 M KCl solution and stored in 3 M KCl at 4 °C.

### Hydrogen peroxide determination

The Pierce Quantitative Peroxide Assay Kit (Aqueous) (ThermoFisher Scientific) was used to quantify peroxide according to manufacturer’s instructions. Briefly, the working reagent (WR) was prepared by mixing 1 volume of Reagent A with 100 volumes of Reagent B, with at least 200 µl prepared for each sample to be assayed. 10 volumes of the WR were added to 1 volume of sample (typically 200 µl WR to 20 µl sample) into a well of a clear bottomed 96 well plate. The reaction was mixed and incubated for 15-20 minutes, after which a Spectramax M3 plate reader was used to measure the absorbance at 595 nm. Sample peroxide concentration was calculated by comparison to a standard curve (dilutions of 30% peroxide) performed the same day.

### Quantification of electrochemical peroxide generation rate

Using experimentally-obtained data for peroxide concentration and charge associated with its electrochemical generation, a characteristic linear relationship between applied charge and peroxide quantity was established for both stirring and non-stirring conditions (**Supplementary Fig. 5**). The efficiency of oxygen reduction to peroxide by the supplied charge was determined by comparing the actual peroxide level to the theoretical yield. The charge was converted to moles of electrons using Faraday’s constant (96485 C mol^-1^) and accounting for two electrons required to produce one hydrogen peroxide molecule. The oxygen reduction efficiency was found to be constant at 0.59 regardless of stirring conditions (**Supplementary Fig. 5**).

### Quantification of cell peroxide consumption rates

Cells were grown overnight as above. To quantify consumption of peroxide, T7Express *E. coli* with the pOxyRS-LacZ-laa plasmid were diluted to optical densities (measured at 600 nm) of 0.025, 0.05, and 0.1, each with 100 µM peroxide in 3 mL of LB medium in 15 mL culture tubes in a 37 °C incubator at 250 rpm. 20 µl aliquots were assayed at each timepoint, and peroxide levels were compared to a standard curve using the Pierce Quantification Peroxide Assay Kit.

For a similar quantification of peroxide consumption from electrode-immobilized cells, the bacteria were first assembled onto gold-coated wafer chips (150 mm^2^) as described by Methods. After rinsing superfluous bacteria from the chips, each was submerged in 2 mL M9 medium supplemented with antibiotics and 100 µM peroxide. Additionally, a negative control containing a sterile gold chip in the solution and a positive control containing suspended cells at an optical density (600 nm) of 0.025 were included. All samples were incubated statically at 37 °C for 2 h, during which 20 µl aliquots were assayed for peroxide concentration at regular timepoints. At the end of the timecourse, the cell-immobilized chips were fixed with paraformaldehyde, followed by fluorescent cell staining with propidium iodide. Image analysis was used to quantify the exact electrode dimensions based on photographs and on-chip cell densities based on fluorescence imaging of the chip surfaces. The rate of peroxide accumulation for each sample was determined by normalizing the assay-measured number of peroxide molecules in solution to cell number.

### Measurement of growth effects of peroxide

To quantify cell growth in the presence of peroxide, *E. coli* T7Express cells with the pOxyRS-sfGFP-laa and pBla-GBP3-AIDA plasmids were cultured overnight as described above. The cells were diluted in the M9 media with antibiotics to an OD600 of 0.025. Either solution-based or electrochemically-generated peroxide was added to the cells at different concentrations, as indicated, in triplicates. A Bioscreen C machine (Growth Curves USA) was set up at 37 °C, high shaking, with 400 µl per well, and recorded OD600 measurements every 15 minutes. The Growth Rates program^49^ was used to calculate lag and doubling time.

### Colony Forming Unit (CFU) determination

Treated or untreated cells were diluted at least 1000 times, and 100 µL of these dilutions were plated on LB + 1.5% agar plates with the appropriate antibiotics. After an overnight incubation at 37 °C, colonies were counted and CFU’s were calculated.

### Flow cytometry

Flow cytometry was used to quantify peroxide-induced sfGFP fluorescence intensity, live/dead cell ratios, and quantum dot labeling of cell surfaces. A BD Accuri C6 with an autosample was used to measure sfGFP and live/dead cell numbersr. Cells were diluted in PBS to achieve less than 10,000 events per second. Reported values are the mean fluorescence values in the FL-1 channel (green) or FL-3 (red).

Quantum dot-labeling of cells was determined using a BD Canto II by first establishing a threshold for unlabeled cells using the 530/30 filter and then comparing above-threshold counts between GBP3- and + strains once mixed with quantum dots. All data is supported by at least 20,000 events and consistently gated for forward and side scatter across samples.

### Miller assay

The Miller assay was performed according to standard protocols. Briefly, cells were lysed with chloroform and sodium dodecyl sulfate (SDS) to release β-galactosidase. The substrate ONPG was added and cleaved by β-galactosidase into a yellow molecule, *o-*nitrophenol. Absorbance of sample sets at 600, 550, and 420 nm was quantified by either a Spectramax M3 or BioTek Synergy plate reader. Absorbance at 600 nm was measured from 250 μL of culture sample prior to assay preparation and the absorbance at 420 and 550 nm were measured from 200 μL of the sample after assaying for β-galactosidase activity. Miller Units were calculated as per the standard protocol.

### Electrochemical PAP measurement

PAP was detected electrochemically through cyclic voltammetry (CV), chronoamperometry (CA), and differential pulse voltammetry (DPV). All measurements were performed with a CHI Instruments 700-series (CH instruments, Inc.) or VMP3 (BioLogic Science Instruments) electrochemical analyzer using gold working electrode (2 or 3 mm diameter, CH Instruments, Inc.), a coiled platinum counter electrode with working area larger than that of the working electrode (BASi), and an Ag/AgCl reference electrode (BASi). CVs were run from −0.15 to 0.3 V at a scan rate of 50 mV/s. The current at the reduction peak was used to measure PAP concentration. DPV was run from -0.16 to 0.36 V, at 2 mV step increments, with a 50 mV amplitude, 0.5 s pulse width, and a 0.5 s pulse period. The DPV peak height was calculated either automatically or manually for calibration to PAP concentration.

### General electroinduction setup

The same electrochemical setup used for peroxide generation was used for induction of electrogenetic cells, and was placed inside of a mini incubator set at 37 °C. Cells were added to a final OD600 of 0.025 in the working electrode vial, or were pre-assembled onto the working electrode itself. The electroinduction solution was M9 medium with the appropriate antibiotics. Peroxide was generated via chronoamperometry as indicated above, with voltage application for a specified duration (*eg* 300 s). For planktonic cells, the entire volume was then pipetted into a 15 mL culture tube, which was placed in a 37 °C shaking incubator at 250 rpm, from which samples were removed at indicated time intervals for measurement. For electrode-assembled cells, the working electrode 15 mL culture tube with fresh media for shaking incubation at 37 °C and 250 rpm. In experiments that electrochemically probed β-galactosidase induction, 5 mg/mL PAPG was added prior to incubation. Cells were assayed as indicated.

### Electroinduction setup with co-cultures

A general electroinduction setup was used for electroinduction of cocultures. The electrogenetic Router cells were initially inoculated to a final OD600 of 0.025 in the working electrode solution for planktonic cocultures or pre-assembled directly on the working electrode, which was submerged in the co-culture. In double-culture experiments, the Verifier cells were co-inoculated to a final OD600 of 0.1. In triple-culture experiments, Verifier and Actuator cells were inoculated to a final OD600 of 0.075 and 0.025, respectively (**Fig. 6b**). Chronoamperometric electroinduction and analysis were performed as in the general protocol above. After electroinduction, the coculture solution and cell-immobilized electrode were transferred to a culture tube for incubation, with 5 mg/mL PAPG added prior to incubation. At indicated time intervals, the cultures were sampled; aliquots were separated by centrifugation as cell pellets for Miller Assays and supernatants for electrochemical PAP detection and protein secretion analysis.

### AHL quantification

AHL quantification was performed by bioluminescence assay. AHL reporter cells JLD271 pAL105 were grown overnight in LB at 37 °C shaker and 250 rpm with the appropriate antibiotics. The following day, standard AHL solutions (concentration range) were prepared in LB. The reporter cells were diluted 2500x in LB with the appropriate antibiotics. To each, 90 uL of diluted reporter cells and 10 µl of the of the standard dilutions were mixed in a 5 mL culture tube and prepared in duplicate. Experimental conditioned media samples were prepared similarly after sterile-filtering and diluting between 5 and 10x to maintain a linear assay range. Culture tubes with reporter cells and conditioned media samples were incubated at 30 °C and 250 rpm shaking for 3 hours. Luminescence was measured with a GloMax®-Multi Jr (Promega, Madison, WI, USA). AHL concentration of each sample was calculated using the standard curve.

### Electrode chip fabrication

Gold electrodes were prepared by cutting gold-coated silicon wafers (Ted Pella) or as purchased pre-cut chips (Platypus Technologies). Gold coating was 50 – 100 nm in thickness with a 5 nm sublayer of chromium or titanium. Alternatively, gold electrode arrays were patterned onto silicon wafers. First, metal deposition was performed on standard 4 inch silicon wafers using a Denton thermal evaporator (Denton Vacuum LLC, Moorestown, NJ, USA), with metal deposition rates at 2–3 Å/s. Specifically, a 50 nm chromium adhesion layer was evaporated first, followed by 200 nm gold. Next, photolithography steps utilized direct writing of photoresist via a DWL66fs laser writer (Heidelberg Instruments, Heidelberg, Germany), guided by a laser exposure map designed in AutoCAD (Autodesk, San Rafael, CA, USA).

Photoresist spin-coating and development steps were performed using an EVG120 Automated resist processing system (EV Group, Sankt Florian am Inn, Austria). The patterned wafer was post-processed by etching, photoresist stripping, and cutting individual electrodes with a DAD dicing saw (DISCO, Tokyo, Japan).

### Protein structure determination

Protein modeling and structural predictions were performed using the Phyre2 web portal^50^. This approach referenced the 4MEE crystal structure^51^ available through the RCSB Protein Data Bank (rcsb.org)^52^ in order to generate a homology model of GBP3-His-AIDAc protein used in this work.

### Protein secretion analysis

The extracellular diffusion of target proteins DsRedExpress2 and GMCSF due to TolAIII expression was evaluated by collecting conditioned media aliquots at designated timepoints of TolAIII cultures. The supernatants of aliquots were recovered after centrifugation (6000 g, 2 min) to separate bacteria and subdivided fluorescence analysis (stored chilled) or immunoassaying (stored frozen). The relative fluorescence intensity of samples was measured using a BioTek Synergy platereader to determine DsRedExpress2 levels. The His-tagged GMCSF peptide was quantified using a His Tag ELISA Detection Kit (GenScript) according to the provided protocol and 5-10 fold dilutions of frozen samples.

### Nanoparticle labeling of bacterial surfaces

For quantum dot labeling, CdSe/ZnS core-shell type quantum dots (Sigma Aldrich, carboxylic acid functionalized, fluorescence λ_em_ 520 nm) were diluted 100-fold into bacterial cultures (OD at 600 nm = 1) resuspended in PBS at a 10X dilution. After incubating 1 h at room temperature, the cell cultures were rinsed twice prior to flow cytometry analysis for quantum dot fluorescence. For gold nanoparticle labeling, 20 nm gold nanoparticles (AuNPs, Ted Pella, #15705-20) were used to label the gold binding peptide constitutively expressed by *E. coli* cells. Transmission electron microscopy (TEM) was performed after labeling cells with Au NPs. Cells were diluted in a 2:1 mixture of PBS and water to an optical density at 600 nm of 0.28. The cells were then mixed with a thousand-fold excess of gold nanoparticles and incubated for one hour. The samples were prepared onto surface-treated copper grids covered with holey carbon films using a drop-casting method, as described by Dong *et al*^53^. The samples were viewed with a JEOL 2100F transmission electron microscope operated at 200 kV.

### Model and simulations

The peroxide, promotor, and protein expression models were developed using first order reaction kinetics to describe OxyR kinetics and Hill functions to describe OxyR activation of gene expression. Models were implemented using Microsoft Excel when no spatial resolution was needed. Spatial dynamics were implemented in Matlab using the built-in PDESolver to solve the diffusion equation. Details on the model, its parameters, and its implementation in Matlab are available in the **Supplemental Methods** and **Supplemental Table 5**.

### Implementation of modeled spatial dynamics

To understand the spatial dynamics of peroxide generation and consumption at the electrode with the cells either absent, present on the electrodes, or present in solution, the diffusion equation was solved in Matlab’s built in PDESolver function. The box size was 3.5 mm per side with a one millimeter square electrode located in the center of one face of the box. The geometry was generated using the GeometryFromMesh function and 1 micron sized increments. Peroxide was generated at the electrode for 300 s and then stopped, reflecting a typical pulse increment. The overall simulation was run for 3600 s. Depending on the simulation, peroxide consumption was either absent (reflecting no cells present), occurring only on the electrode (reflecting immobilized cells), or in solution (reflecting peroxide pulse into a bulk culture). The equations describing this behavior have been detailed in the **Supplemental Methods**, the parameters that used for the peroxide diffusion, generation and consumption are noted in **Supplemental Tables 5**, and experimental derivations of simulation parameters are described in **Supplemental Table 6**.

## Author Contributions

JL Terrell, T Tschirhart, Y Liu, CY Tsao, HC Wu, G Vora, GF Payne, DN Stratis-Cullum, and WE Bentley were involved with the conception and design of the work. JL Terrell, T Tschirhart, K Stephens, R McKay, M Pozo, and HC Wu were involved with engineering strains used in this work. JL Terrell, T Tschirhart, JP Jahnke, K Stephens and H Dong were involved with data acquisition and interpretation. JP Jahnke and MM Hurley were involved with computational kinetic studies and protein modeling, respectively. JL Terrell, T Tschirhart, JP Jahnke, GF Payne, DN Stratis-Cullum, and WE Bentley were involved with writing and documentation of the work.

## Acknowledgements

Partial support of this work was provided by DTRA (HDTRA1-19-0021), NSF (DMREF #1435957, ECCS#1807604, CBET#1805274), the National Institutes of Health (R21EB024102), and the Office of Naval Research (N0001417WX01318, N0001418WX01042). This work was also supported by the Assistant Secretary of Defense for Research and Engineering (ASD(R&E)) through the Applied Research for Advancement of S&T Priorities Synthetic Biology for Military Environments program.

## Data Availability

The datasets that support the findings of this study are available from the corresponding author upon request.

## Code Availability

The Matlab code for the models used in this study is available from the corresponding author upon request.

## Supplementary Information

### Supplementary Methods

#### Cloning of peroxide-inducible genetic constructs

To construct a plasmid with a hydrogen peroxide-inducible circuit, the plasmid pOxyRS-sfgfp-laa was first engineered. First, the empty vector pBR322 was amplified with primers pBR322-1 and pBR322-2. The OxyR-pOxyS region was isolated from a boiled colony of *E. coli* MG1655 using OxyRS-1 and OxyRS-2. sfGFPlaa was isolated from pTGX(ref) using sfGFPlaa-1 and sfGFPlaa-2. After DpnI treatment, fragments were ligated by Gibson Assembly. This plasmid product, pOxy-sfGFPlaaV1, linearized with primers pOxyGFP-1 and pOxyGFP-2, and *proD*, amplified with primers proD-1 and proD-2, were ligated by Gibson Assembly to yield the final construct, pOxy-sfGFPlaa. Other genes of interest, *lacZ* and *lasI*, were substituted into the POxy construct, to create variants including and excluding the *ssRA* degradation tag (pOxy-LacZ, pOxy-LacZlaa, pOxy-LasI, and pOxy-LasIlaa). Here, pOxy-sfGFPlaa was linearized by PCR using PoxyAssem-R and PoxyAssem-F (to exclude the ssrA *laa* tag) or ssrA-F (to include the ssrA *laa* tag). Likewise, *lacZ* was amplified from pET-LacZ (**Supplementary Table 1**) using PoxyLacZ-F and PoxyLacZ-R (for *lacZ* product) or LacZ-ssrA-R (for *lacZlaa* product); *lasI* was amplified from pLasI (**Supplementary Table 1**) using PoxyLasI-F and PoxyLasI-R (for *lasI* product) or LasI-ssrA-R (for *lasIlaa* product). Each amplified gene was ligated into a corresponding pOxy or pOxy-laa backbone by Gibson Assembly to yield the final constructs and transformed into *E. coli* strain NEB10β (**Supplementary Table 2**). Sequences of listed PCR primers are denoted in **Supplementary Table 3**.

#### Cloning of surface display genetic constructs

To develop a plasmid construct for surface display of a gold-binding peptide, pGBP3 was engineered. First, the *AIDA* fusion gene with its preceding signal peptide was amplified from pET-Venus-AIDA (**Supplementary Table 1**) using SigPep_pBla-ovhg-F and AIDA_pBla-ovhg-R primers. The host plasmid, pBla was linearized by PCR using pBla_5end-R and pBla_3end-F and the template pGB-FimH (**Supplementary Table 1**) to open an insertion site at the bla promoter. The signal peptide-AIDA fragment was inserted into pBla by Gibson Assembly to achieve pBla-AIDA for constitutive AIDA expression under the bla promoter. *Gbp^3^* was purchased from IDT with a 5’ extension, TGCATTTGCAGTCGAC, preceding the ATG start codon. As such, GBP3assem-F and GBP3assem-R were used to amplify the gene. Additionally, the pBla-AIDA plasmid was linearized by PCR for GBP3 insertion downstream of the signal peptide using SurfDispIns-R and SurfDispIns-F; the two fragments were ligated by Gibson Assembly to yield pGBP3 for expression of the fusion gene: signal peptide-GBP^3^-linker-AIDA. Variations of pGBP3 were prepared as control surface display constructs. pHis was prepared by plasmid PCR using GSHis6-F and SurfDispIns-R; pLinker was prepared by plasmid PCR using KpnI-Linker-F and SurfDispIns-R. Each PCR was followed by phosphorylation, ligation, and transformation of the DNA product. Final plasmid constructs pGBP3, pHis, and pLinker were cotransformed with pET-LacZ or pOxy plasmids (pOxy-sfGFPlaa, pOxy-LacZlaa, pOxy-LasIlaa) in *E. coli* strains NEB10β, BL21(DE3), or T7 Express (**Supplementary Table 2**). Sequences of listed PCR primers are denoted in **Supplementary Table 3**.

#### Cloning of acyl homoserine lactone reporter genetic constructs

Acyl homoserine lactone responsive genetic constructs were developed using pAHL_reporter_Red_Green as a plasmid backbone (**Supplementary Table 1**). To develop pAHL-LacZ, the sfGFP reporter gene was first replaced with a counterpart containing additional restriction endonuclease recognition sequences. *Sfgfp* was modified by PCR to include a SpeI site at the 5’ flank and both SacI and BamHI sites at the 3’ flank using primers sfGFP_SpeI-F and sfGFP_BamHISacI-R. Both the plasmid backbone and sfgfp insert were digested with SpeI and SacI and ligated to yield pAHL_reporter_Red_Green*. Next, LacZ was amplified using LacZ_SpeI-F and LacZ_BamHI-R, then inserted into the pAHL_reporter_Red_Green* backbone by digesting each component with SpeI and BamHI and ligation to yield pAHL-LacZ. Alternatively, the pAHL-GMCSF construct was prepared by Gibson Assembly. Assembled components included the GMCSF-TolAIII construct, amplified by PCR from pRM102 (**Supplementary Table 1)** with pAHL-TolAIII-F and pAHL-TolAIII-R, and vector pAHL_reporter_Red_Green vector, linearized by PCR using pAHLassem-F and pAHLassem-R primers. pAHL-LacZ was transformed into an LW7 host and pAHL-GMCSF was transformed into a NEB10-β host (**Supplementary Table 2**). Sequences of listed PCR primers are denoted in **Supplementary Table 3**.

#### Model of hydrogen peroxide consumption by *E. coli*

Bacterial consumption of hydrogen peroxide was modeled as a pair of differential equations (**Equations (S1-2)**). As shown in **Supplementary Fig. 1b**, under the conditions studied here, *E. coli* cells are the dominant consumer of hydrogen peroxide (H_2_O_2_) and, in the absence of bacteria, the peroxide concentration is nearly constant. Across the range of cell densities examined here, peroxide consumption by the cells can be modeled as a simple first order equation (**Equation (S1)**). Because of the time scale of the peroxide dissipation, it is also necessary to consider the continued growth and division of the bacteria (**Equation (S2)**). Rate constants *k_1_* and *k_double_* are reported in **Supplementary Table 5.** *k_1_* is obtained from fitting the peroxide concentration over time in **Supplementary Fig. 1b**, using the approximation of 10^9^ cells per 1 OD600 unit. In **Supplementary Fig. 6(c)**, the doubling time ranged from ca. 45 to 60 min and the rate constant (*k_double_*) was taken from the higher end of this range.

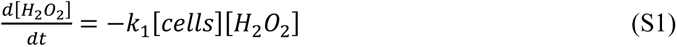

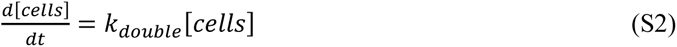

#### Model of hydrogen peroxide diffusion from an electrode

Matlab was used to simulate the spatial distribution of hydrogen peroxide during its electrochemical generation. The diffusion of hydrogen peroxide (H_2_O_2_) from the electrode was modeled with the diffusion equation **Equation (S3)**, taking 8.8 x 10^-6^ cm^2^ · s^-1^ as the diffusion constant of hydrogen peroxide in water^1^.

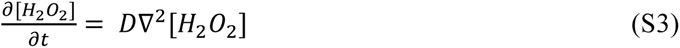

For the flux at the boundary, no hydrogen peroxide is generated or consumed except at the gold electrode surface. The flux of hydrogen peroxide generated at the electrodes using non-stirring conditions was taken from the measurements of hydrogen peroxide production in our system (**Supplementary Fig. 5**), using the smallest electrode size implemented in this work (1 mm^2^) as the surface area unit; using a different electrode size would give very similar results. For conditions where cells are present in solution or at the electrode, an additional consumption term is included (**Equation (S1)**). Model parameters and their derivations from experiments are listed in **Supplementary Table 6**.

#### Model of OxyR-regulated protein expression

The behavior of the OxyR promoter was modeled as a set of first order reactions where OxyR is oxidized by peroxide and returns to the reduced state at a rate proportional to the amount of oxidized OxyR. The oxidized (OxyR(o)) and reduced (OxyR(r)) fractions sum to a constant value, normalized to 1 in this model. This results in a single differential equation to describe the OxyR oxidation state, **Equation (S4)**. Rate constants are reported in **Supplementary Figure 5** and are consistent with literature^2^. Since the same promoter system regulates all protein expression systems modeled here, the parameters are kept constant across all protein models. OxyR(o) levels are shown in **Supplementary Fig. 1c** with 10 min timepoints and 25 µM data shown in **Fig. 2b** and **c**, respectively.

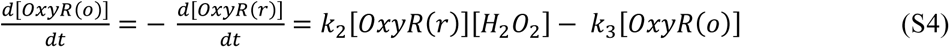

To model OxyR oxidation as a consequence of peroxide uptake during its electrochemical generation (**Fig. 3a-b**), the consumption rate of peroxide by the cells rather than the bulk peroxide concentration was used to calculate the rate of OxyR oxidation. This can be related to the bulk concentration model using **Equation (S1)**, resulting in **Equation (S5)**.

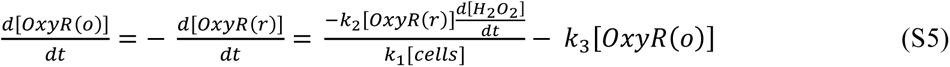

The expression of three different proteins under the OxyR promoter was modeled (β-gal, sfGFP, and LasI). For the activity of the OxyR promoter itself, a Hill equation was used, modified by a proportionality constant to adjust for differences between the assays used to experimentally quantify the individual protein levels (*k_7_*). The specific equations for the proteins are listed below, with the rate constants summarized in **Supplementary Table 5**.

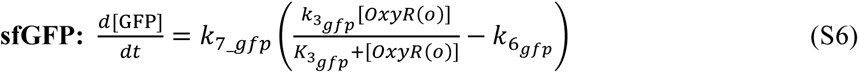

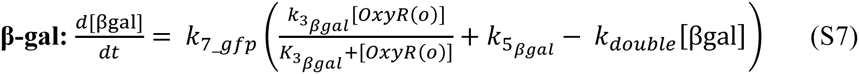

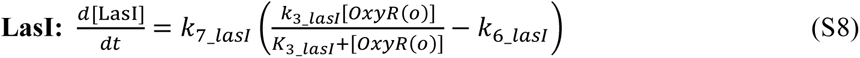

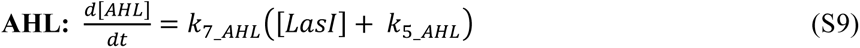

The constants *k_3_* and *K_3_* were fit to the β-gal expression levels at different hydrogen peroxide amounts (**Supplementary Fig. 2**). For the β-gal, its ssrA degradation tag was ineffective in accelerating protein degradation, leading to an assumption of slow degradation compared to the timescale of the experiment. However, the protein is diluted as cell growth and division occurs, and this effect is included in the model (**Supplementary Fig. 2a**). Additionally, there is a low level of leaky expression of the β-gal even without any addition of peroxide, and a parameter fit from the β-gal activity without peroxide is included to reflect this (*k_5_*). Both the experimental data and modeled fits for dose-response between 0 and 100 µM peroxide and a timecourse of 180 min are plotted in **Supplementary Fig 2b**. The experimental data and resulting fit were each saturation-normalized (to the maximum value in Miller Units) and plotted in **Fig. 2b-c**.

For sfGFP and LasI, *K_3_* was taken from the β-gal expression since the promoter behavior should be similar across all three proteins; *K_3_GFP_* was adjusted slightly. For protein degradation, sfGFP and LasI both have active ssrA degradation tags while that of β-gal is ineffective. A zero order reaction was used to represent the degradation kinetics expected at high protein concentrations, where the degradation machinery is expected to be saturated (**Supplementary Fig. 2a**). This constant, *k_6_*, was fit from the experimental AHL levels produced by LasI at 25 µM peroxide induction (**Supplementary Fig. 2e**).

For sfGFP, the expected fluorescence intensities over time are displayed in arbitrary units that are equivalent to the median fluorescence intensities of experimental flow cytometry data (**Supplementary Figs. 2d and 3**). Further, the expected fluorescent population fraction (% of total population) was calculated based on these trends using an error function.^3^ For a normally distributed function with mean (µ) and standard deviation (σ), the fraction of the function above a given threshold (τ) is given by **Equation (S10)**.

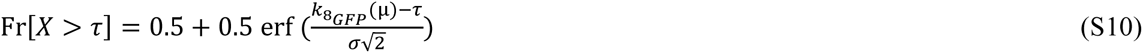

The logarithm of fluorescence in experimental flow cytometry data was approximately normally distributed with a standard deviation of 0.35 in a base 10 logarithm (**Supplementary Fig. 3**). τ was fit to the data to scale for both the overall fluorescence intensity and the measurement threshold. The µ values were taken from modeled sfGFP expression profiles (**Supplementary Fig. 2c**) at 45 min, corresponding to the time where experimental data was also collected (**Supplementary Fig. 3**). The experimental data and resulting fit were each saturation-normalized (to 100 %) and plotted in **Fig. 2b**.

In the case of the LasI, the assay used in these experiments does not directly reflect the LasI concentration but rather the AHL produced by the LasI. This was modeled as a first order reaction in protein concentration assuming a large excess of the AHL precursor with the reaction constant k_7_AHL_ subsuming both the LasI concentration (arbitrarily scaled using k_7_LasI_) and its reaction rate; this constant is scaled based on the amount of AHL produced when 25 µM hydrogen peroxide is used to induce LasI expression (**Supplementary Fig. 2e**) Additionally, baseline AHL is included as k_5_AHL_, which is attributed to noise between basal AHL formation and bioluminescence in the detection assay (**Supplementary Fig. 2f**). The experimental data and fits for LasI and AHL are reported in **Supplementary Figs. 2e**; additionally, AHL data were normalized to the maximum concentration value and plotted in **Fig. 2c**.

## Supplementary Figures

**Supplementary Fig. 1.**
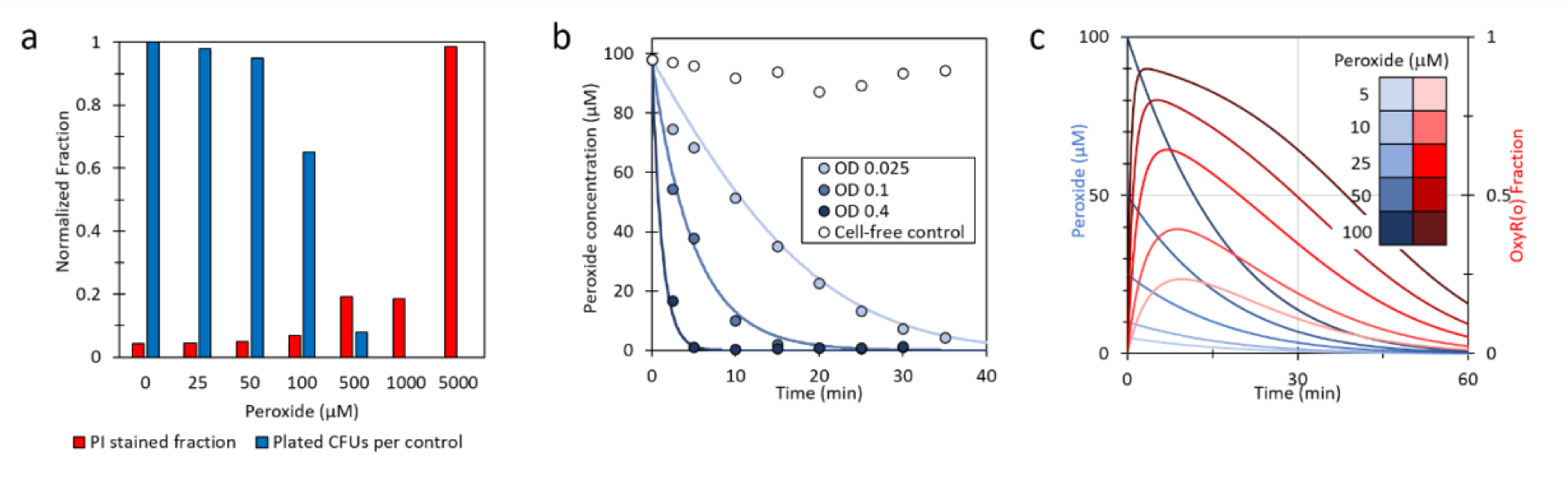
Evaluation of *E. coli* interactions with hydrogen peroxide. (a) Bacterial viability after peroxide dose exposure, measured for non-viable cells by counting a propidium iodide stained cell fraction and for viable cells by counting colony forming units. (b) Peroxide depletion data for cultures at varying cell densities (0.025, 0.1, and 0.4 OD600) shown as experimental measurements (circles) and modeled behavior (lines). (c) Modeled extracellular peroxide concentration and resultant intracellular fraction of OxyR(o) over time for peroxide concentrations between 0 to 100 µM.

**Supplementary Fig. 2.**
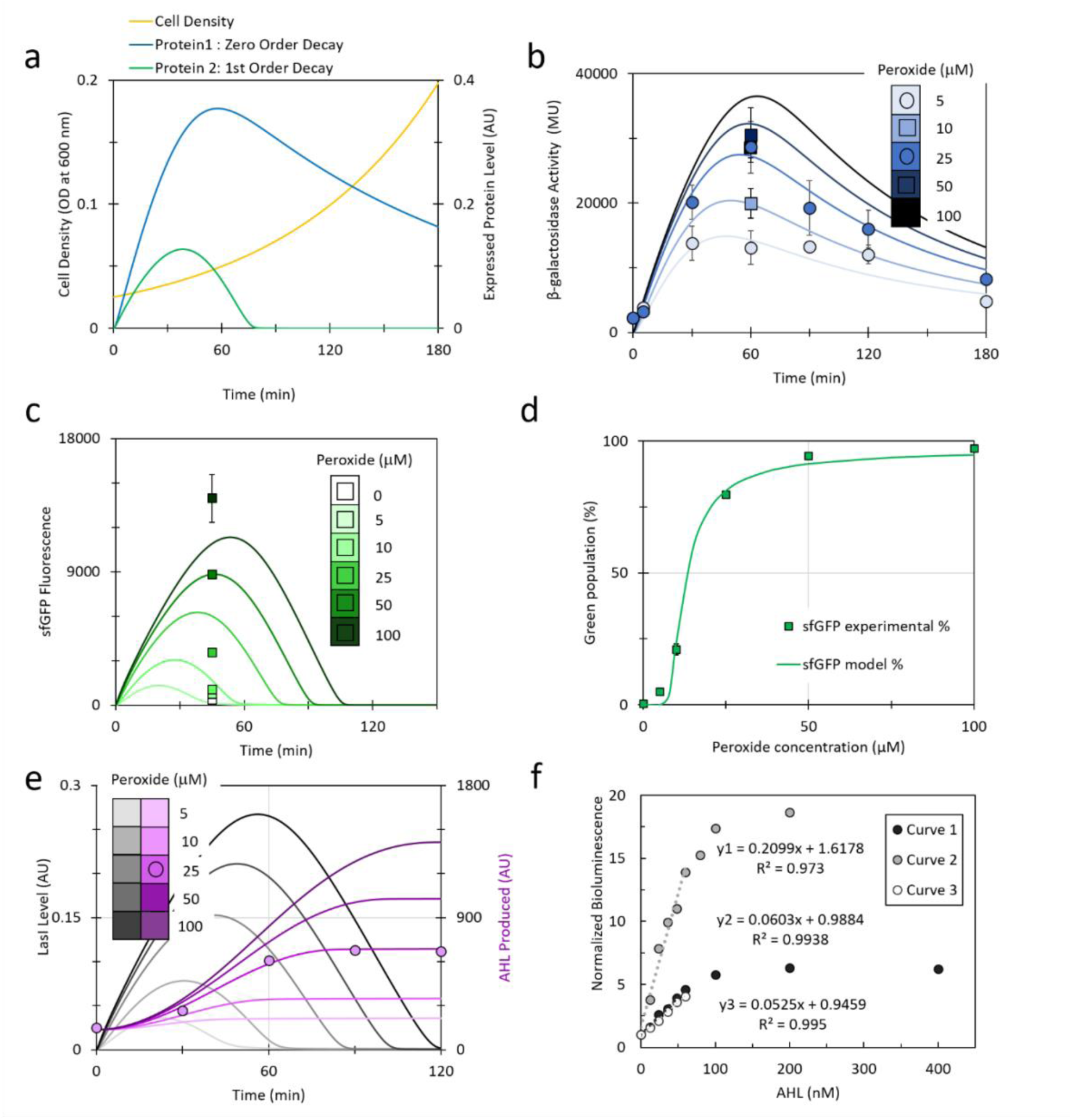
Modeled experimental data for hydrogen peroxide-regulated protein expression and activity. **(a)** Timecourse of cell density and 25 µM-induced expression levels of a protein exhibiting either zero-order decay due to cell division or first-order decay influenced by degradation tag. Modeled OxyR-regulated protein expression timecourses across a range of 0 to 100 µM peroxide adjusted to experimental data for **(b)** β-galactosidase, **(c)** sfGFP fluorescence, **(d)** percentage of sfGFP-expressing cells in a population, **(e)** LasI level and LasI-dependent acyl-homoserine lactone (AHL). **(f)** Standard curves for AHL concentrations between 0 and 400 nM, measured by bioluminescence assay. For plots (b-e), color-coded legends indicate peroxide condition for models (solid lines); square data points indicate experimental dose-response results and circles indicate experimental timecourse results. Experimental data for (b-d) were obtained in technical triplicate with averages and standard deviation indicated while data for (e) were quantified in duplicate by bioluminescence assay, using Standard Curve #2 in (f).

**Supplementary Fig. 3.**
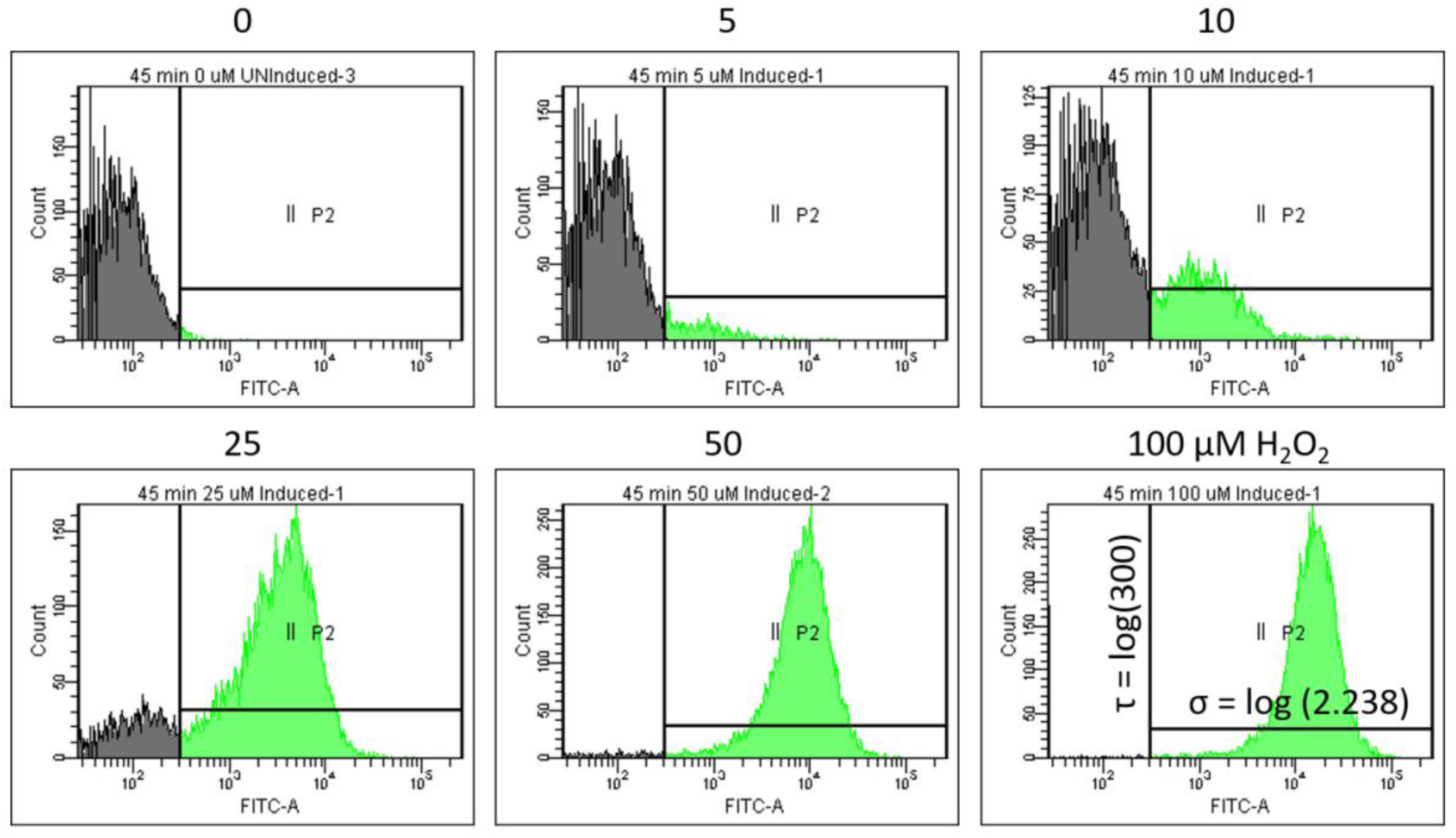
Fluorescence histograms for peroxide-induced sfGFP expression. Experimental flow cytometry data for pOxy-sfGFP cells after 45 min induction with hydrogen peroxide (0 – 100 µM). Population distribution model parameters (τ, σ) were derived from the base 10 logarithm of the fluorescence threshold and standard deviation, given a normal distribution.

**Supplementary Fig. 4.**
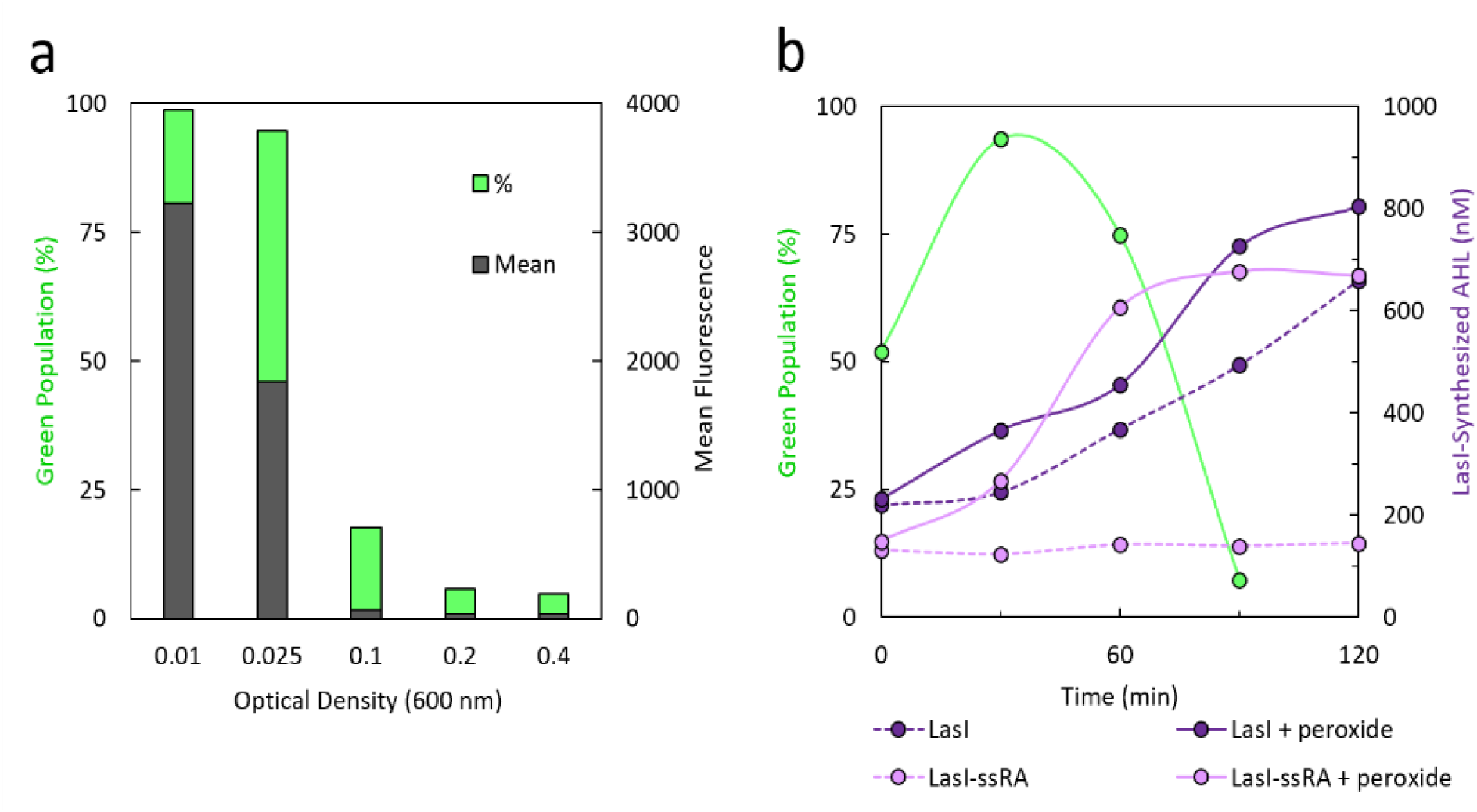
Factors affecting electrogenetic protein expression. **(a)** Influence of cell density on sfGFP fluorescence (as green population % exceeding fluorescence threshold and mean relative fluorescence of population) when induced with 100 µM peroxide. **(b)** Timecourse expression when induced with 25 µM peroxide of sfGFP fluorescence (with ssRA degradation tag) and LasI-synthesized acyl-homoserine lactone (AHL, quantified by Curve 1 (**Supplementary Fig. 2-2**), compared between LasI with and without the ssRA tag inclusion.

**Supplemental Fig. 5.**
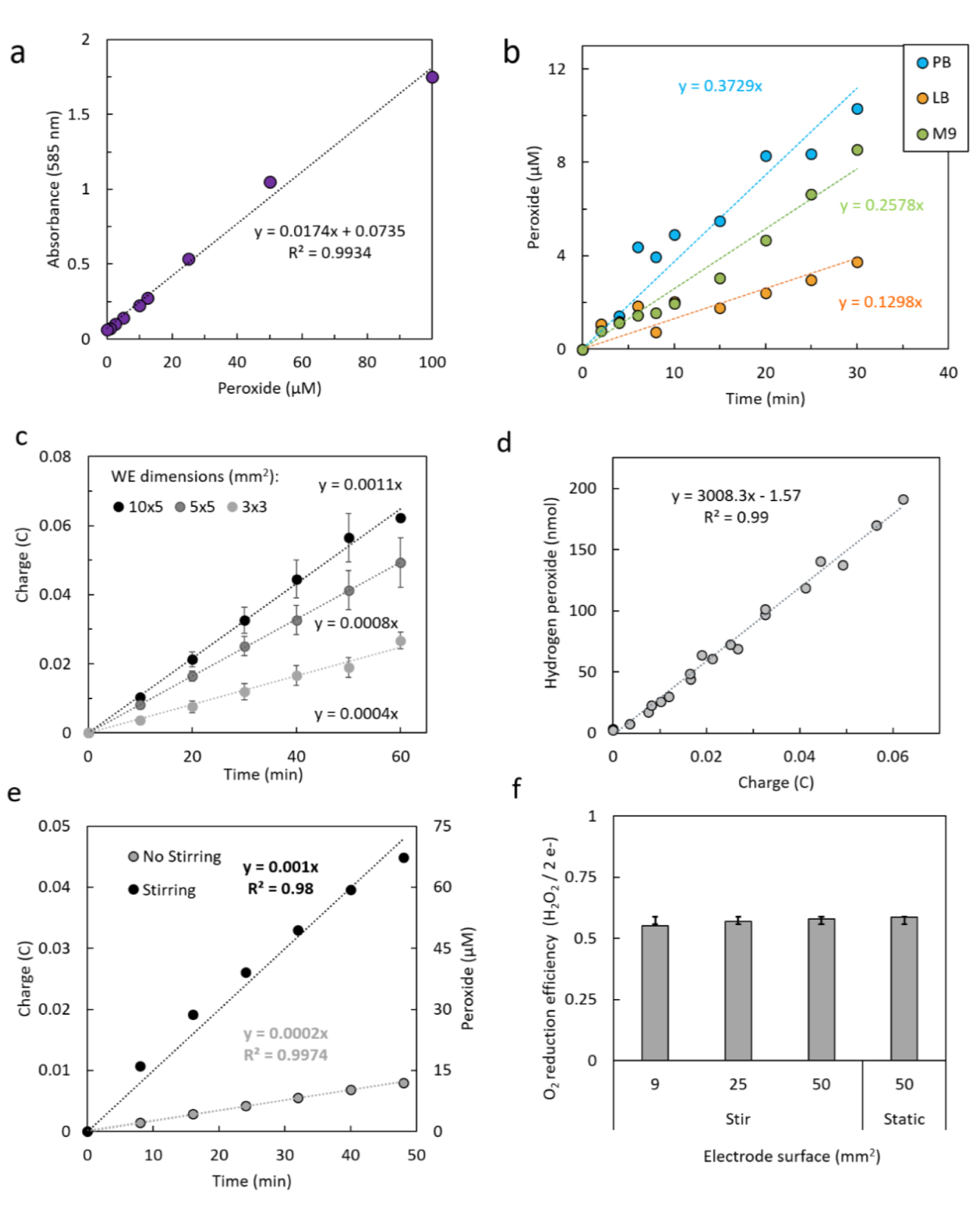
Characterization of electrochemical peroxide generation rate. **(a)** Standard correlation between hydrogen peroxide concentration in M9 medium and spectrophotometric absorbance at 585 nm using a colorimetric peroxide assay. **(b)** Peroxide accumulation over time with constant voltage in various physiologically-compatible solutions: LB (lysogeny broth), M9 (minimal medium), PB (phosphate buffer). All set-ups used a split electrochemical configuration with conductivity across a salt bridge. PB was also tested with electrochemistry occurring in a single chamber (no salt bridge). **(c)** Charge transfer during electrochemical peroxide generation using electrodes of indicated surface area. Inset shows picture of representative wells of Peroxoquant assay of time-course peroxide generation from indicated electrodes. **(d)** Correlation of applied charge to accumulated peroxide molecules based on data in (c) and Figure 2(e). **(e)** Differences in rates of charge transfer and corresponding peroxide accumulation as a function of aeration (stirring or no stirring). **(f)** Experimental stoichiometric efficiency of oxygen reduction to yield hydrogen peroxide for various electrode dimensions and stirring conditions (see **Supplemental Methods** for calculation). Standard deviation of mean efficiency (0.58) shown by error bars. In c, data represents the means of technical triplicates and error bars their standard deviation.

**Supplementary Fig. 6.**
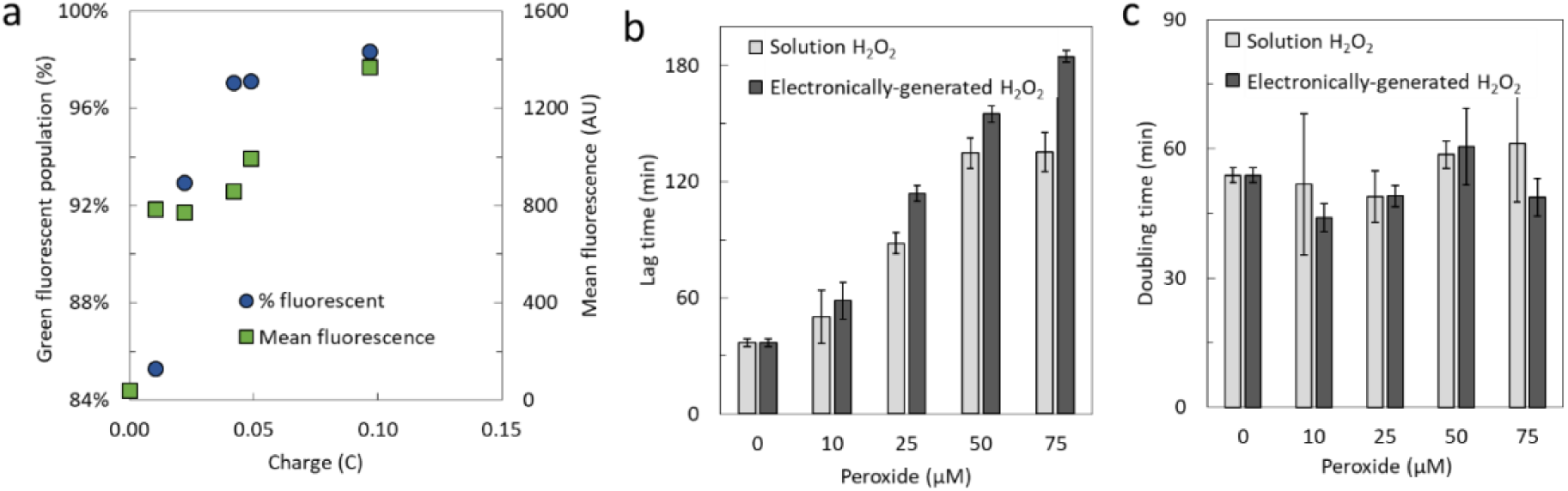
Electrogenetic induction and effects on cell growth. **(a)** sfGFP fluorescence, measured by flow cytometry, of cells induced with the indicated peroxide-generating charges after 45 minutes. **(b)** Lag time growth effect and **(c)** doubling time growth effect of bacterial cultures with either exogenously-added or electrochemically-generated peroxide. Data in b and c calculated using Growth Curves program (Supplementary Methods). Bar graphs depict the means of technical triplicates and error bars represent their standard deviations.

**Supplementary Fig. 7.**
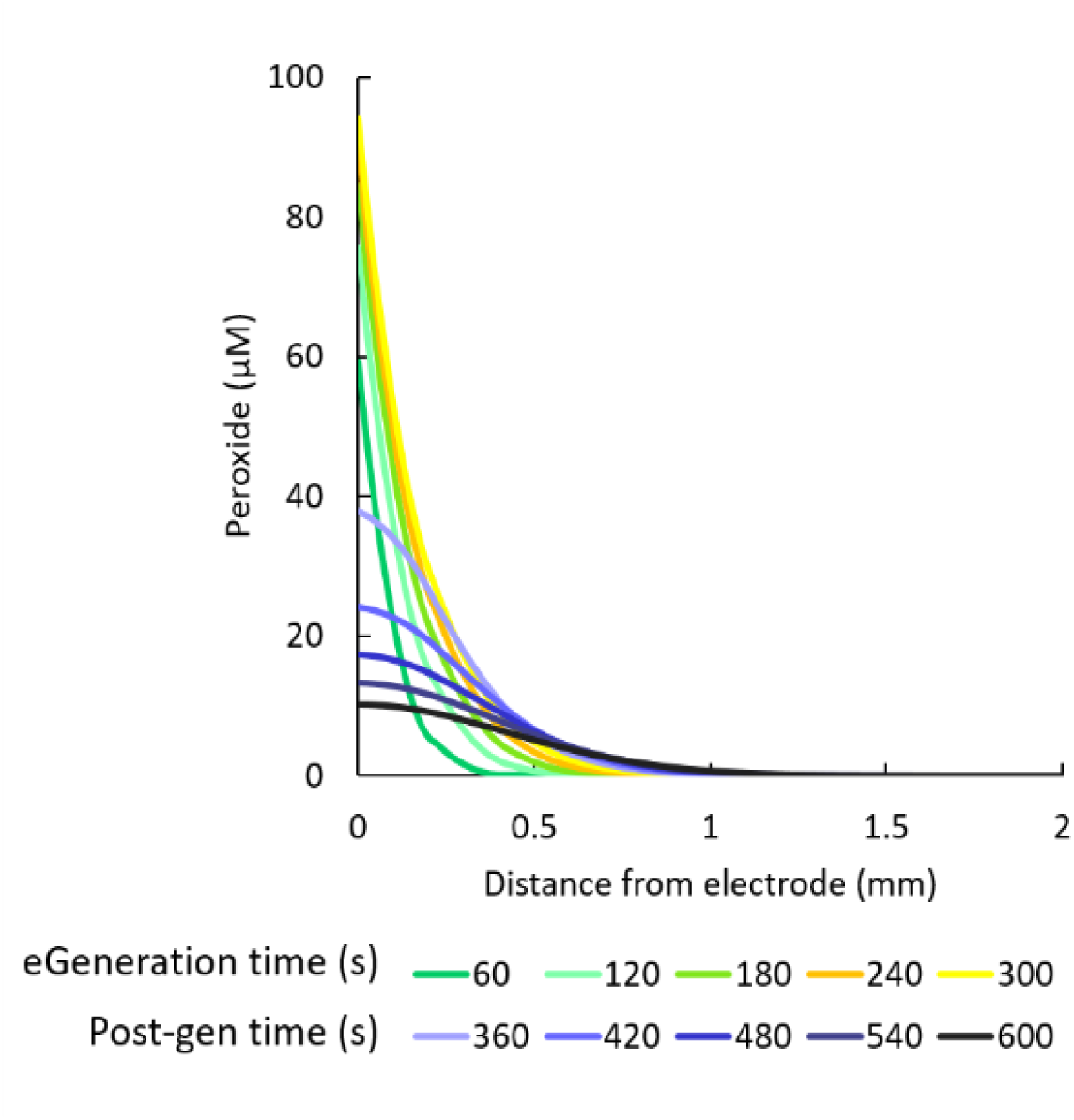
Simulated electro-generated peroxide levels at (yellow/green) and after (blue) specified duration, in systems without the presence of cells.

**Supplementary Fig. 8.**
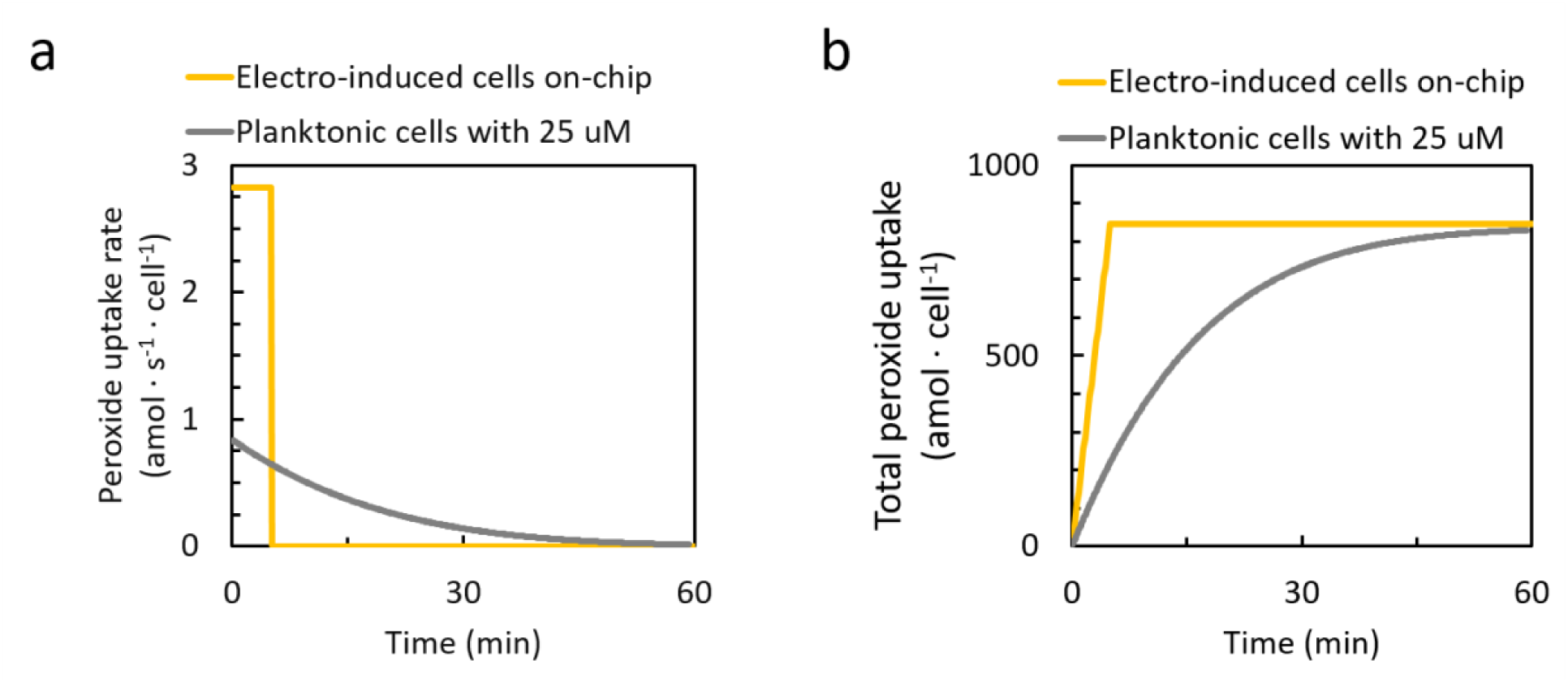
Comparison of peroxide uptake between peroxide-induced planktonic cells and electro-induced electrode-immobilized cells. Per-cell peroxide uptake **(a)** rate and **(b)** cumulative amounts for cells at 0025 OD induced with 25 µM peroxide or cells-on-chip elctro-induced for 300 s.

**Supplementary Fig. 9.**
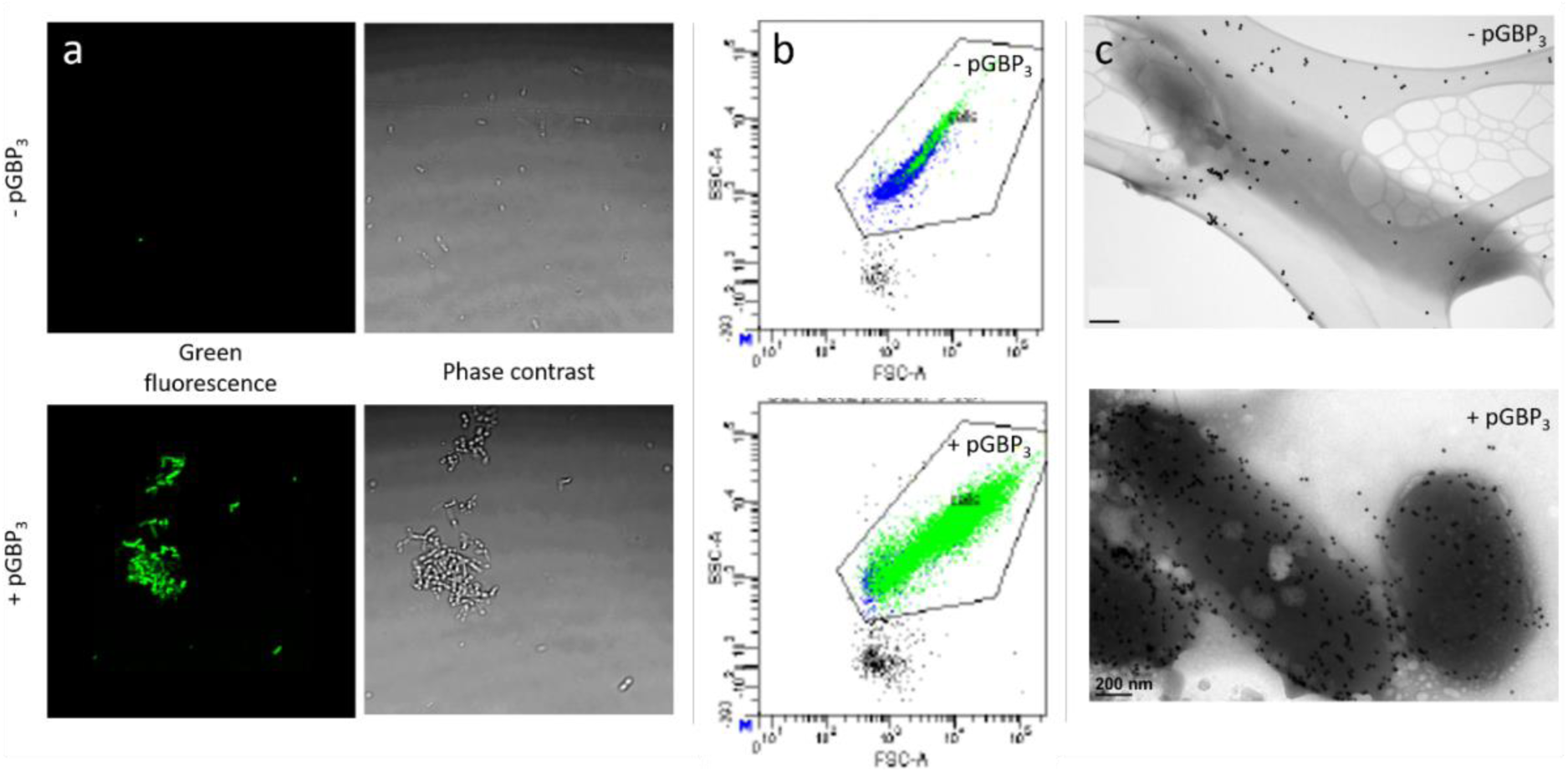
Metal nanoparticle binding to cell surfaces with surface display of GBP^3^-His^6^ peptide. **(a)** Corresponding fluoresence and phase contrast images of *E. coli* cells and CdSe/ZnS quantum dots (QD, em = 520 nm). **(b)** Forward/Side scatter distributions of QD-labeled *E. coli* with the fluorescent-gated population represented by green data points. **(c)** Transmission electron microscopy images of *E. coli* with 20 nm gold nanoparticles. (a-c) Top row shows GBP^3–^ cells. Bottom row shows GBP^3+^ cells.

**Supplementary Fig. 10.**
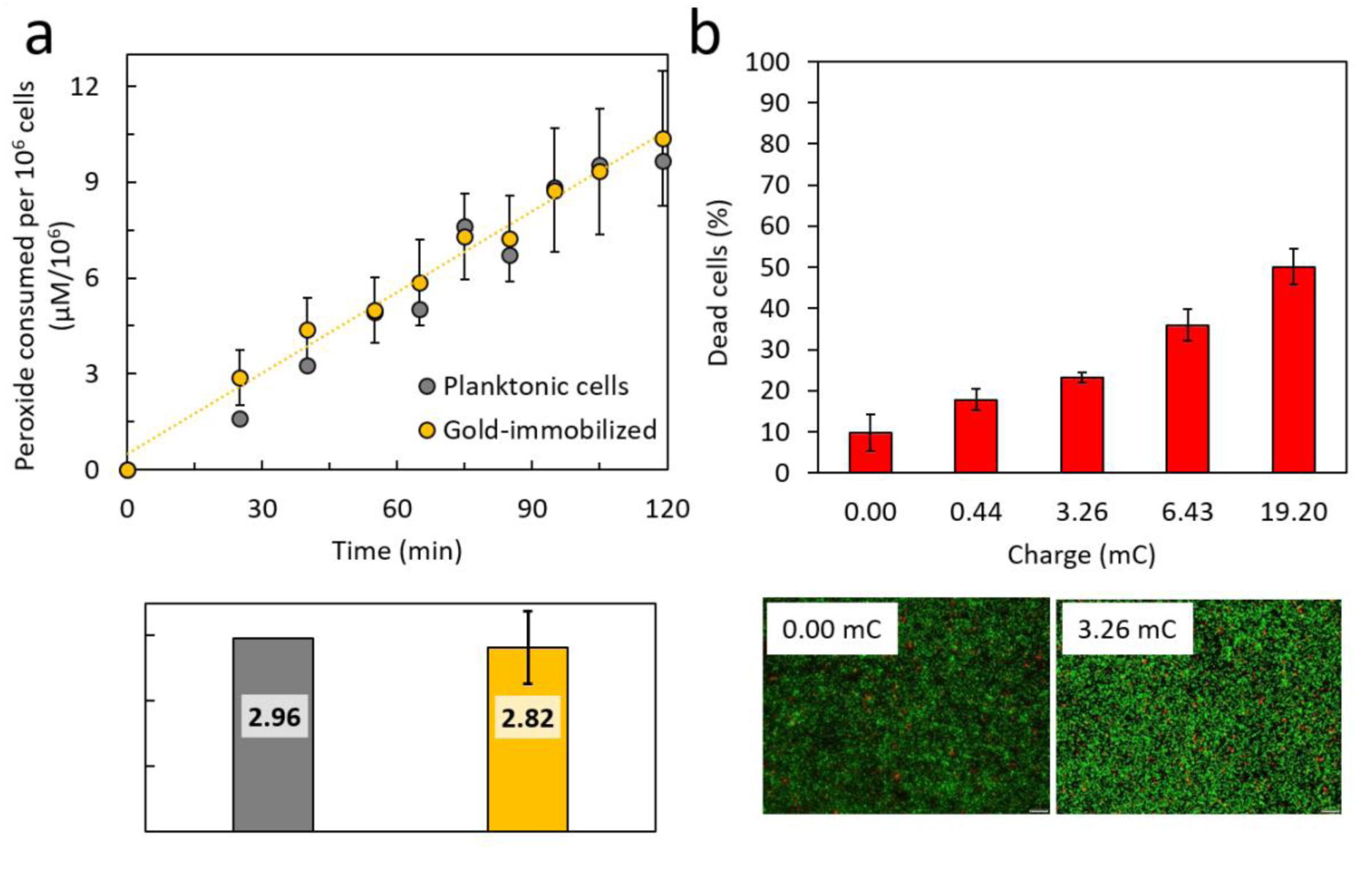
Characteristics of gold-immobilized cells. **(a)** Measured peroxide depletion by electrode-immobilized or planktonic cells, with concentration over time subtracted from the starting level, 100 µM. Bottom plot shows the consumption rate of each normalized to cell number, determined by starting optical density for planktonic cells or image analysis of fluorescently stained immobilized cells. All data for gold-immobilized samples is an average of four biological replicates with standard deviation indicated by error bars. **(b)** Observed cell viability (% dead) of electrode-immobilized cells after exposure to varying electro-inducing charges, based on image analysis of live/dead fluorescence staining on-chip done by ImageJ. Means of at least biological triplicates and the corresponding standard deviations are shown. Bottom pictures show composite red/green fluorescence images of PI + Syto-9 –stained GBP^3^-immobilized cells after electro-induction with the indicated charge or no induction.

**Supplementary Fig. 11.**
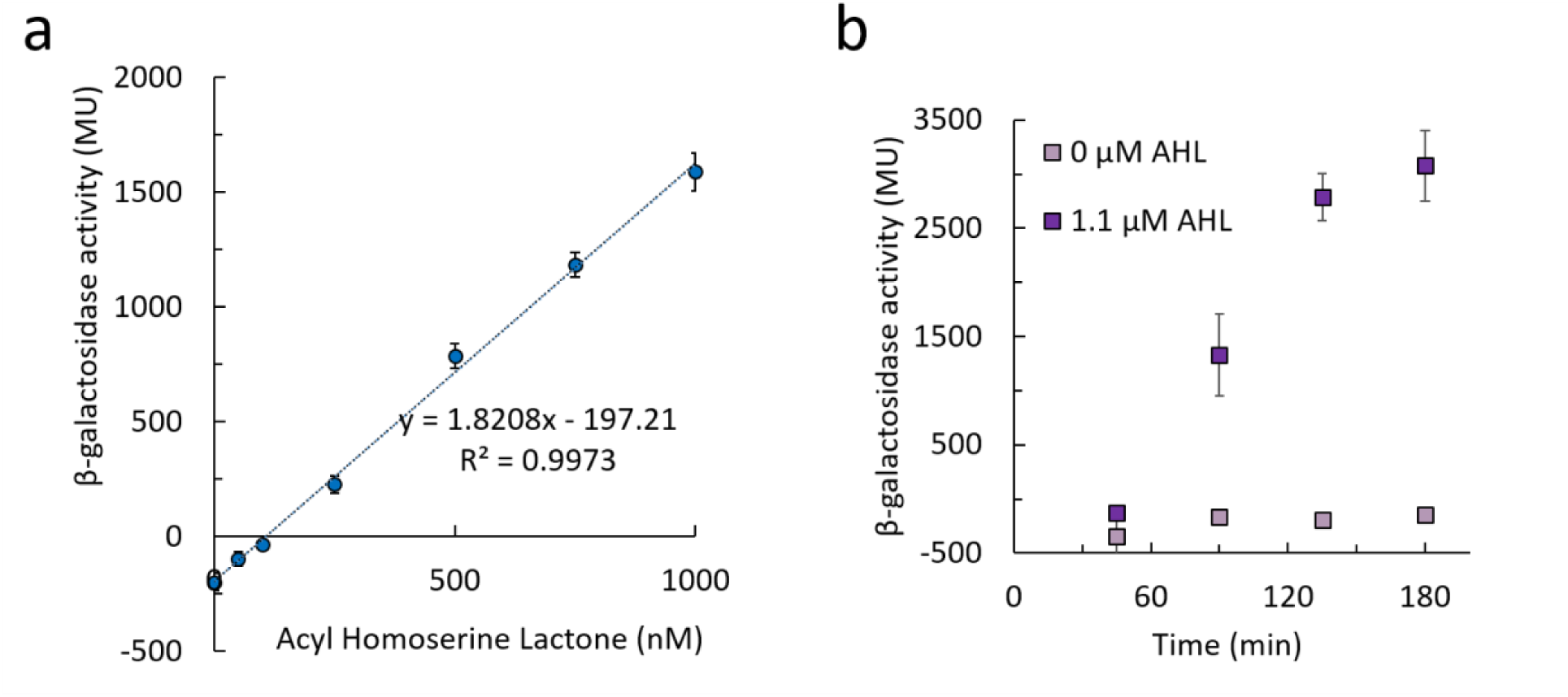
Characterization of LacZ reporter cells in response to acyl homoserine lactone (AHL) induction. For pAHL-LacZ^+^ cells, measurement of β-galactosidase activity (in Miller Units): **(a)** as a response to an AHL dose up to 1.1 µM induction at 90 min and **(b)** as a timecourse without induction (0 µM) or with 1.1 µM AHL introduced at 0 min. All data represent the averages of technical triplicates, with standard deviation reported.

**Supplementary Fig. 12.**
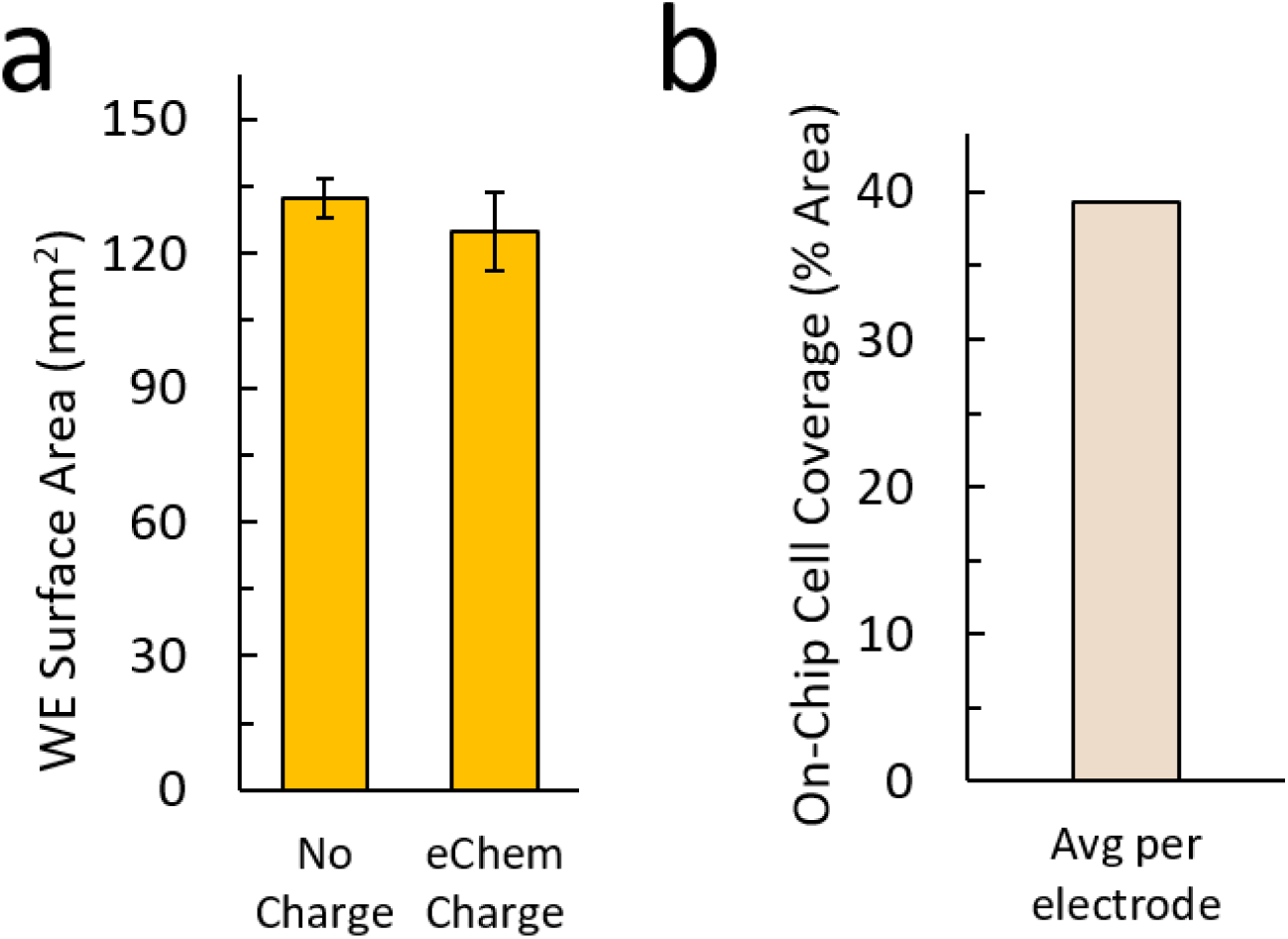
Average surface area of working electrodes **(a)** and cell coverage **(b)** in the experiment shown in Fig. 5d.

**Supplementary Fig. 13.**
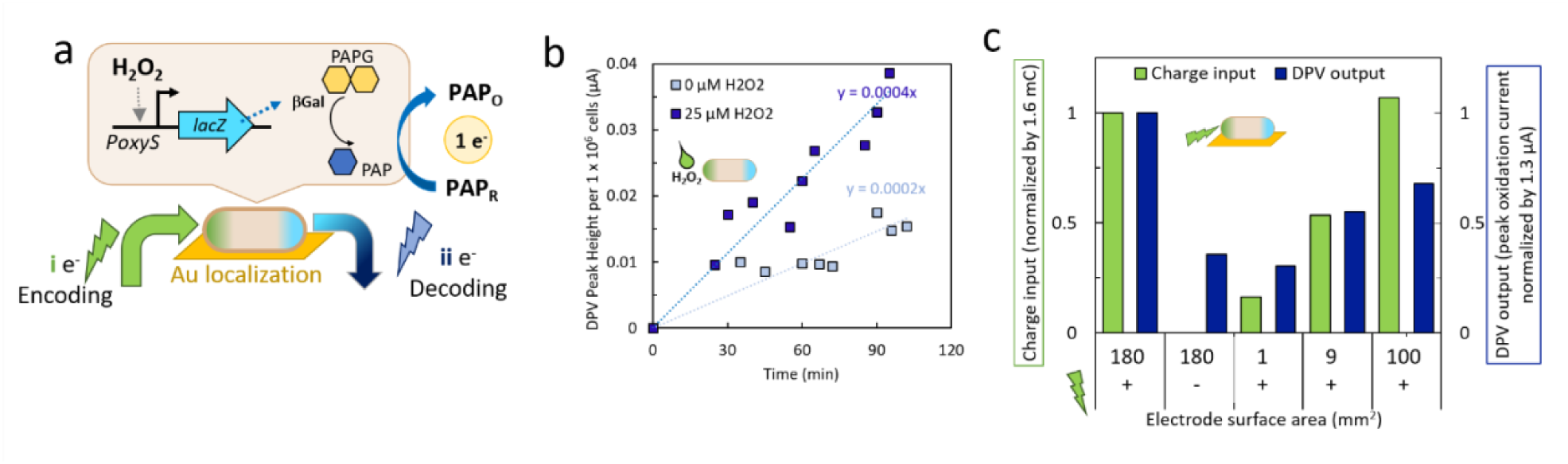
Electronic information flow through a single bi-directionally-connected cell type. **(a)** Schematic of information flow from the electrode, through the cell (peroxide induction of LacZ, and enzymatic PAP generation) and back out to the electrode, where PAP oxidation provides an electronic output. **(b)** Electronic output, quantified by DPV peak current of PAP oxidation, from cells with or without exogenous peroxide induction, measured over time. **(c)** Control-normalized charge input (peroxide formation) and cell electronic output (PAP oxidation peak current), from electrode-immobilized cells. Systems were set up with differing electrode surface areas, as indicated, with other system parameters remaining the same.

**Supplementary Fig. 14.**
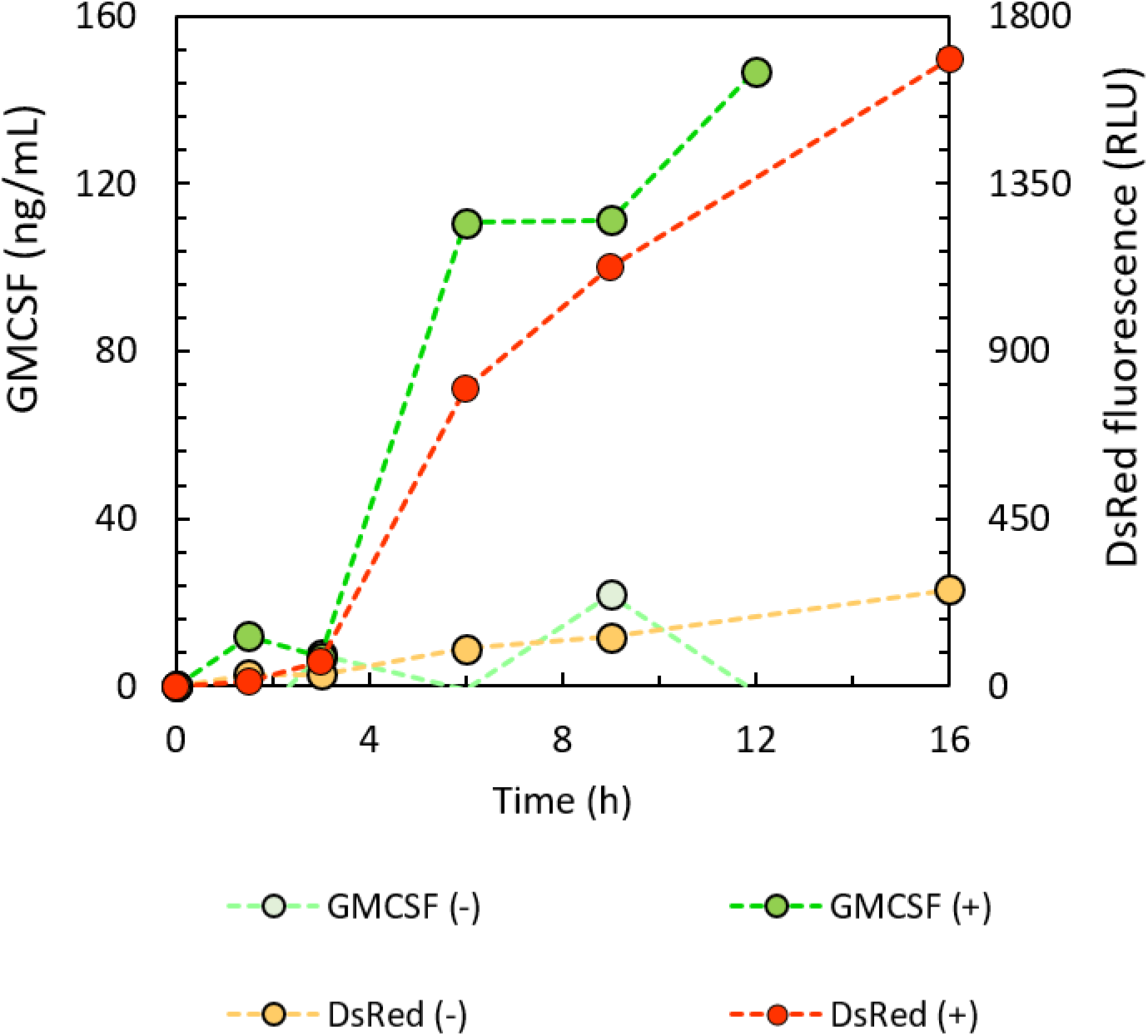
Timecourse levels of extracellular GMCSF and DsRed secreted by Actuator cells either uninduced (-) or induced with 1.1 µM acyl homoserine lactone (+).

**Supplementary Table 1.**
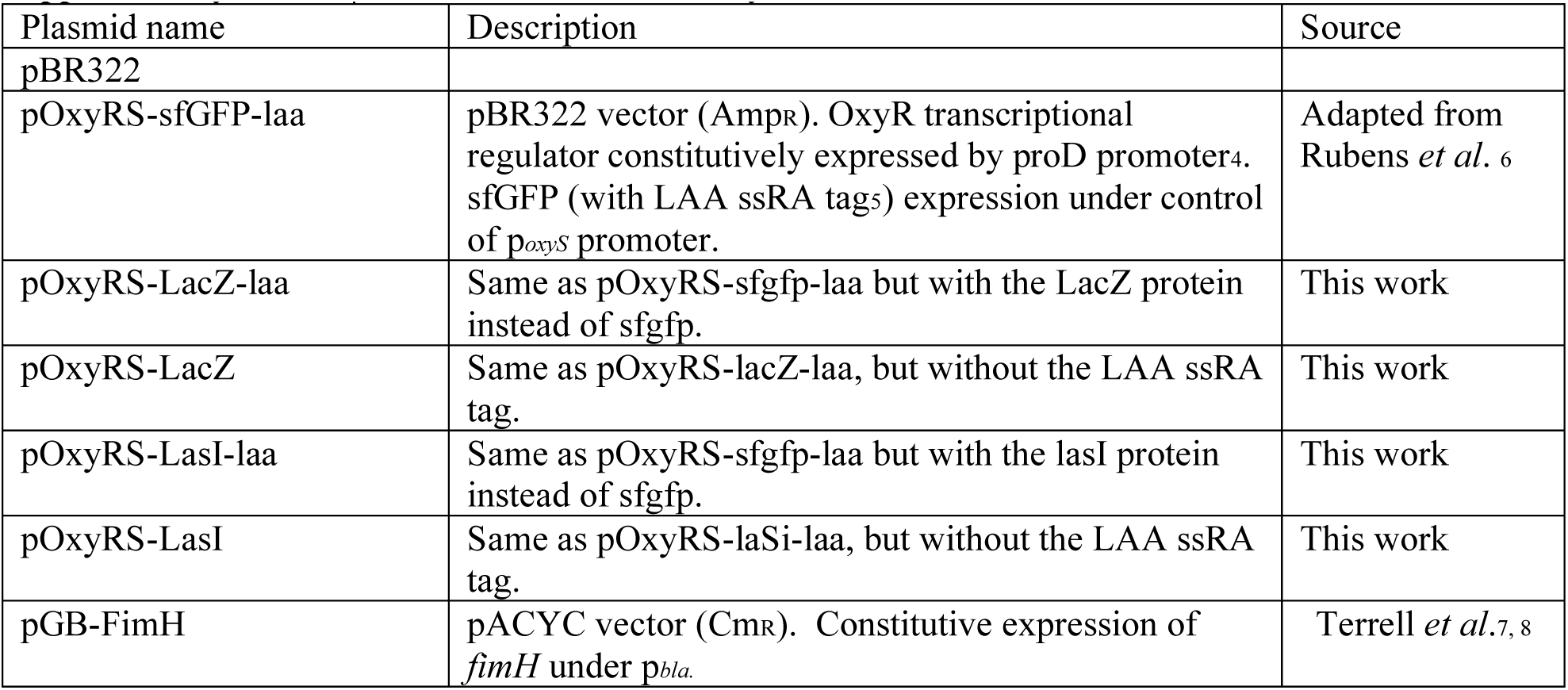
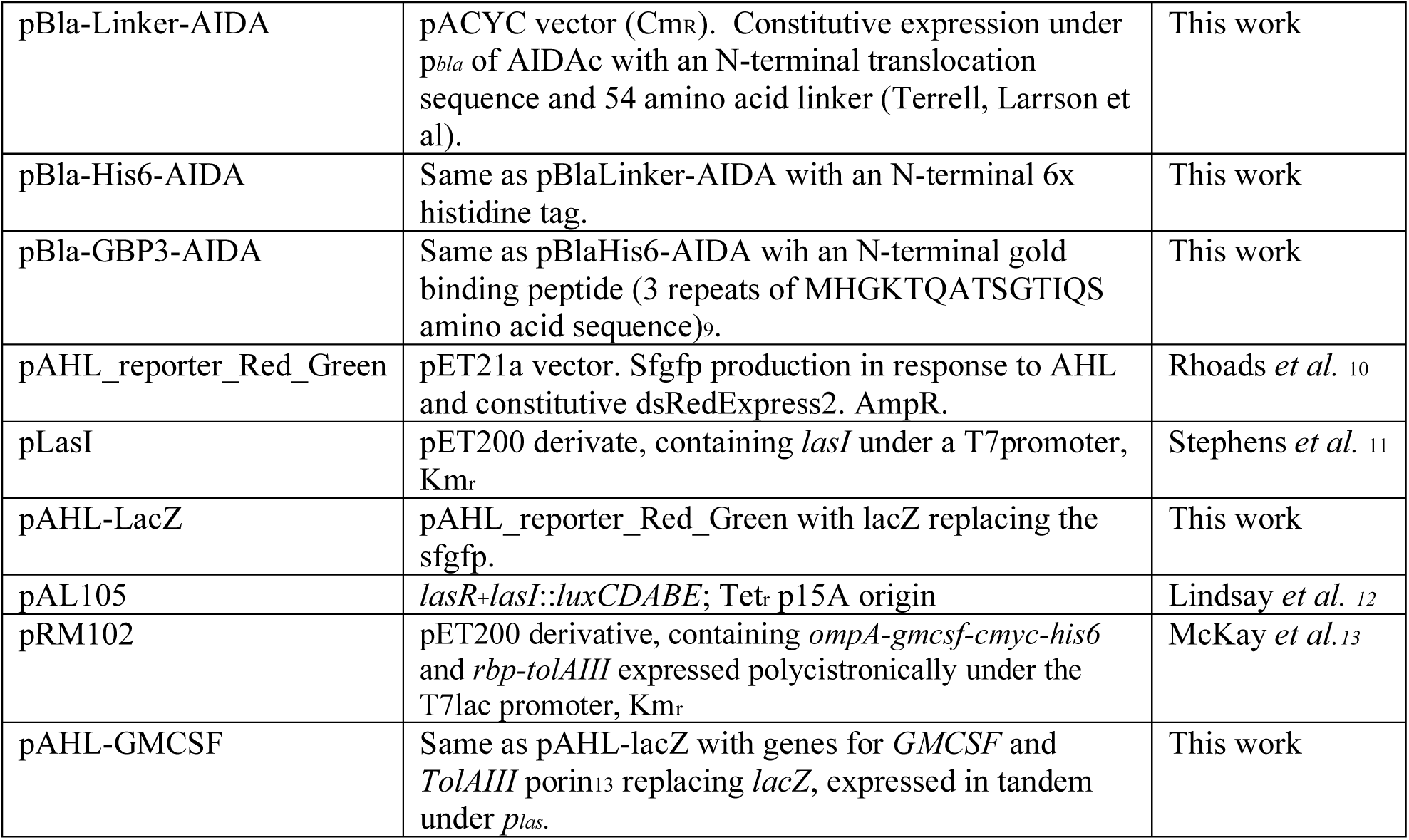
Plasmids used in this study.

**Supplementary Table 2.** Cells used in this study.

| Cell name | Genotype | Source |
| --- | --- | --- |
| NEB10β | $\Delta(ara-leu)$ 7697 *araD139 fhuA $\Delta lacX74 galK16 galE15 e14-$ $\phi 80dlacZ\Delta M15 recA1 relA1 endA1 nupG rpsL$ (Str<sup>R</sup>) *rph spoT1 $\Delta(mrr-hsdRMS-mcrBC)$** | New England Biolabs |
| LW7 | ZK126 $\Delta luxS::Kan$ | Wang *et al.* <sup>14</sup> |
| BL21 (DE3) | *fhuA2 [lon] ompT gal ($\lambda$ DE3) [dcm] $\Delta hsdS$ $\lambda$ DE3 = $\lambda$ *sBamHI* $\Delta EcoRI-B$ *int::(lacI::PlacUV5::T7 gene1) i21 $\Delta nin5$** | New England Biolabs |
| T7Express | *fhuA2 lacZ::T7 gene1 [lon] ompT gal sulA11 R(mcr-73::miniTn10--Tets)2 [dcm] R(zgb-210::Tn10--Tets) endA1 $\Delta(mcrC-mrr)114::IS10$* | New England Biolabs |
| JLD271 | *E. coli* K-12 $\Delta lacX74 sdiA271::Cam$ | Lindsay *et al.* <sup>12</sup> |

**Supplementary Table 3.**
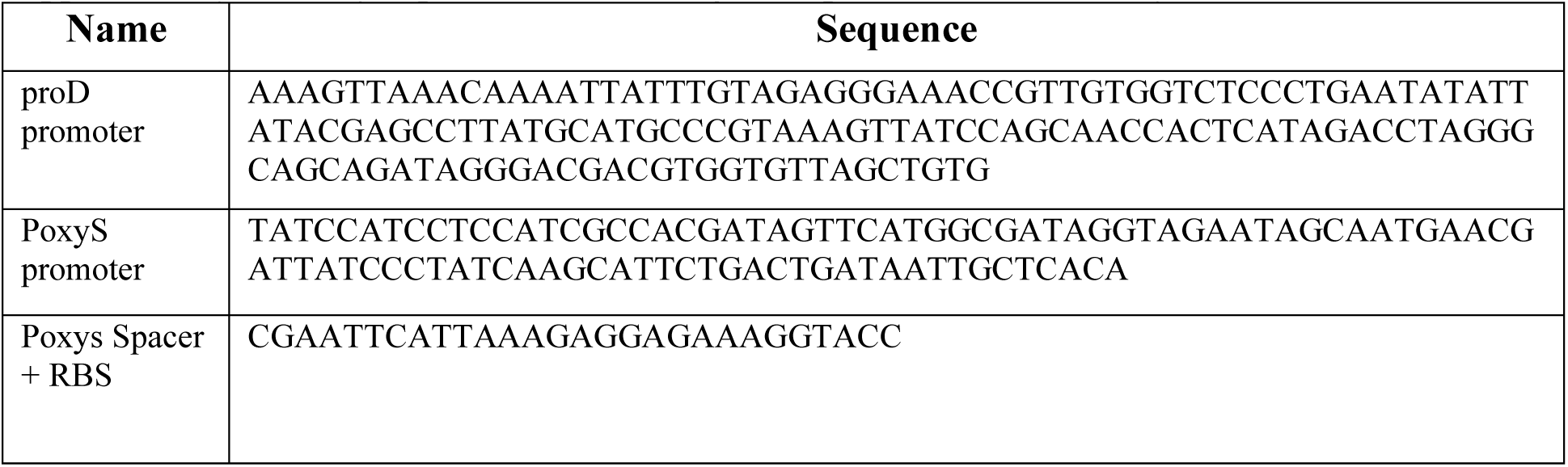
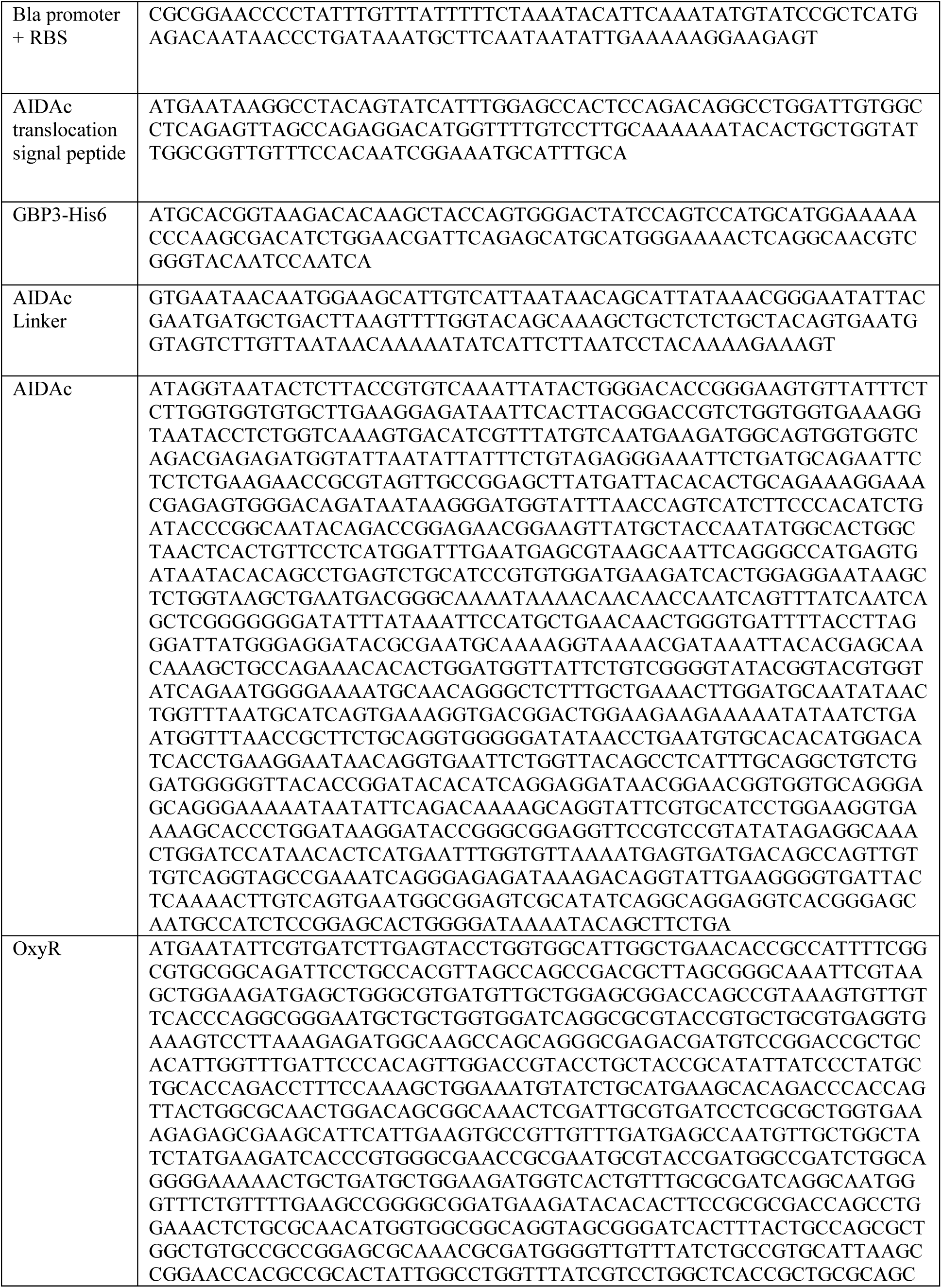
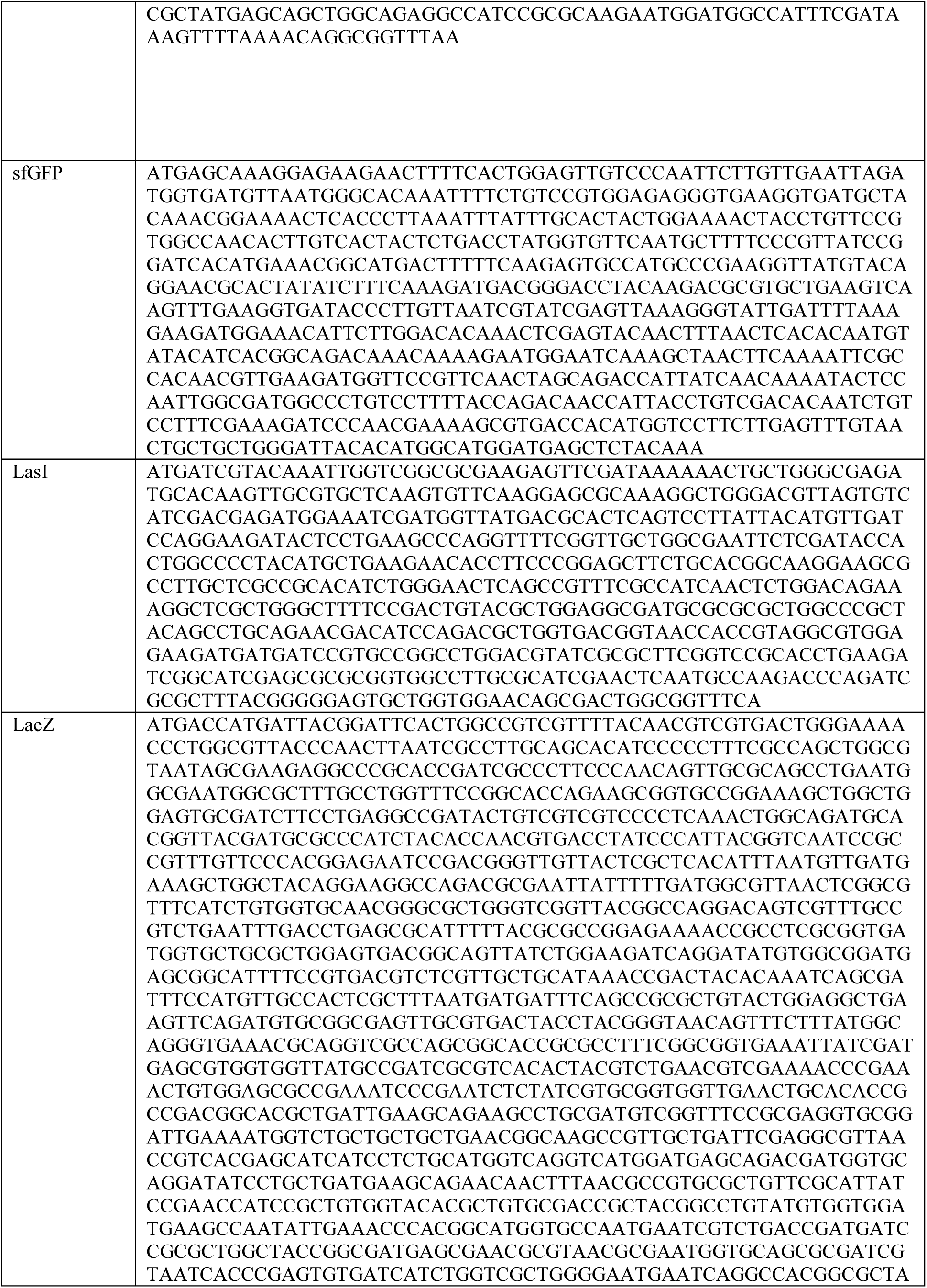
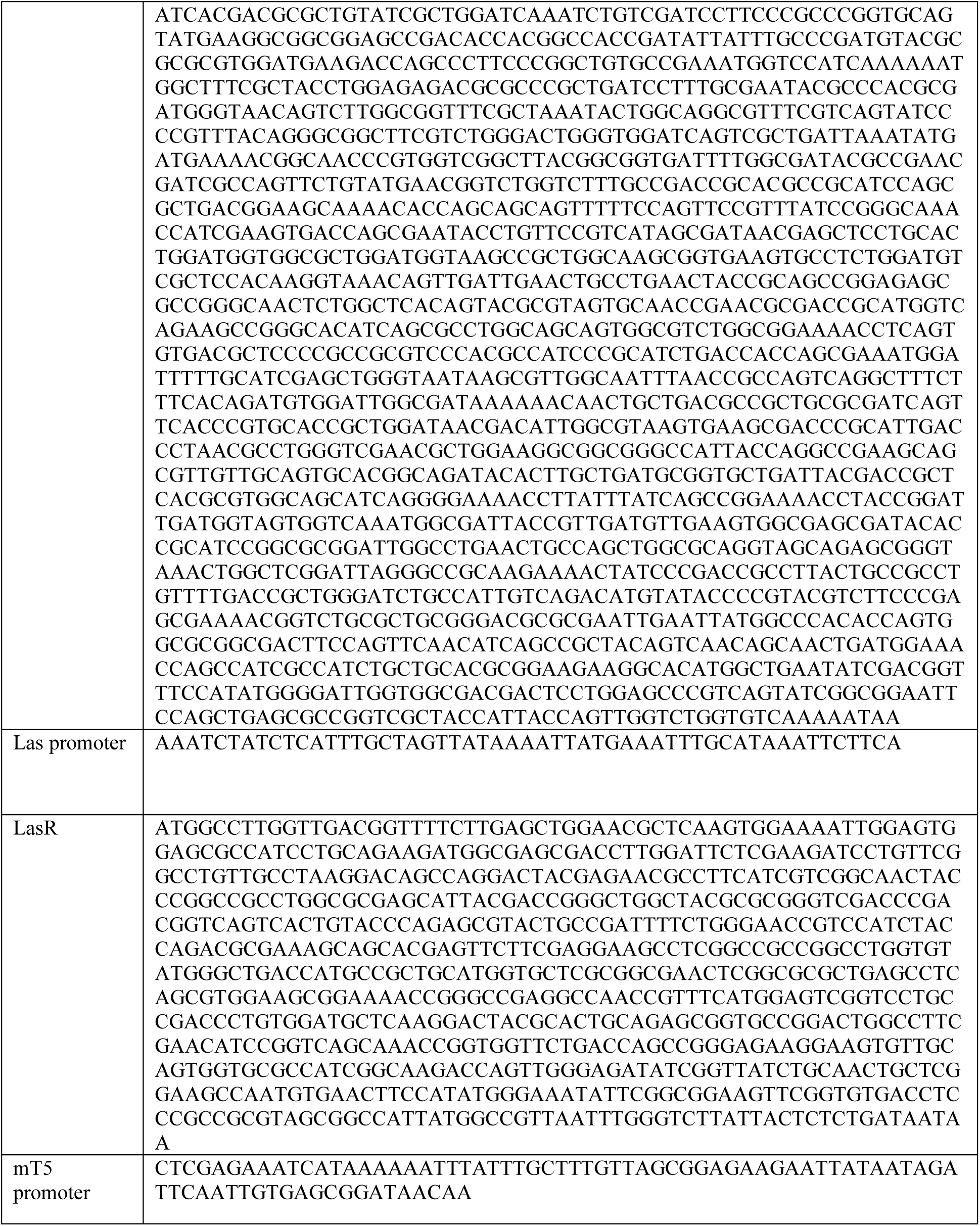
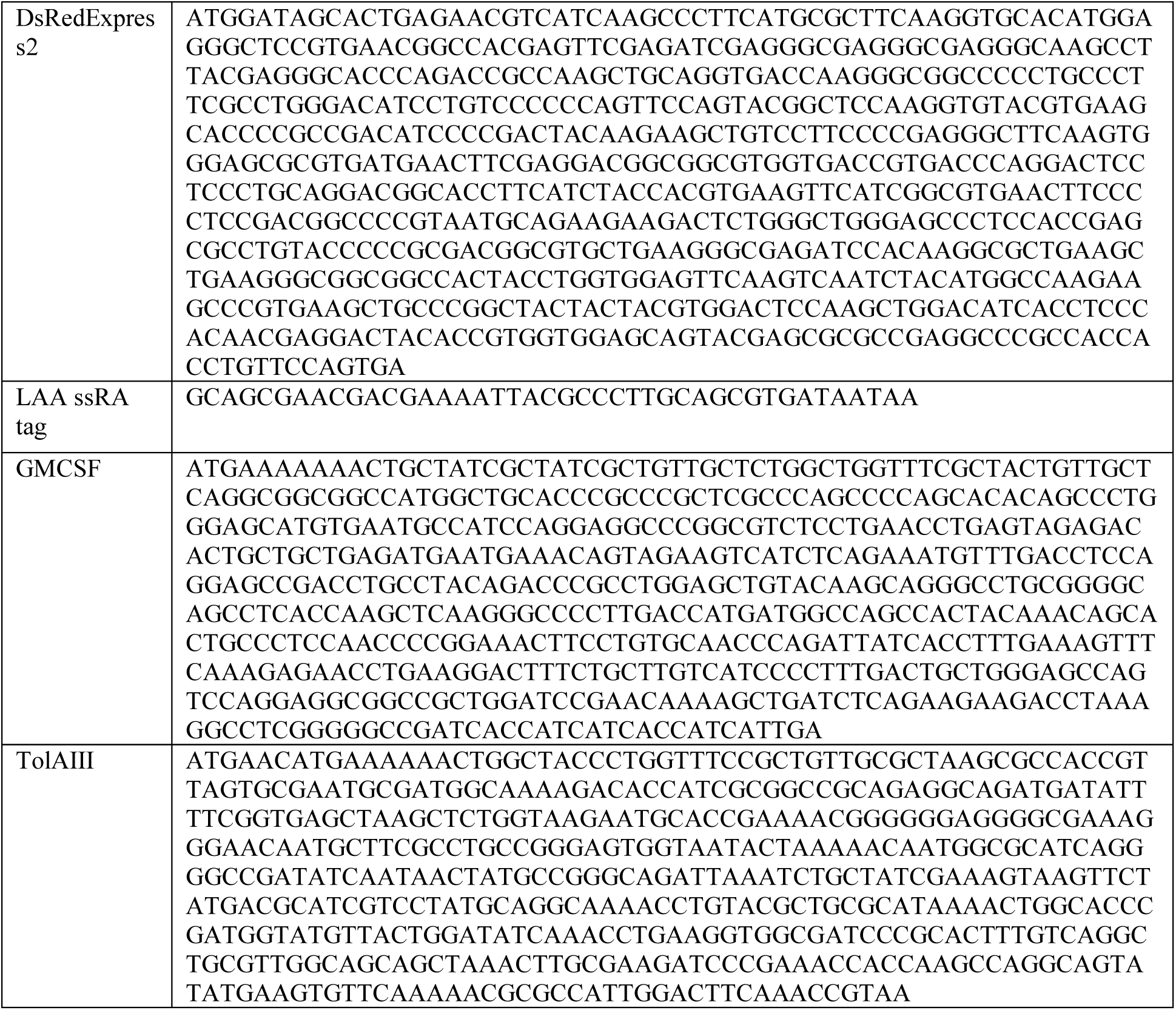
Sequences of relevant genetic parts used in this study.

**Supplementary Table 4.**
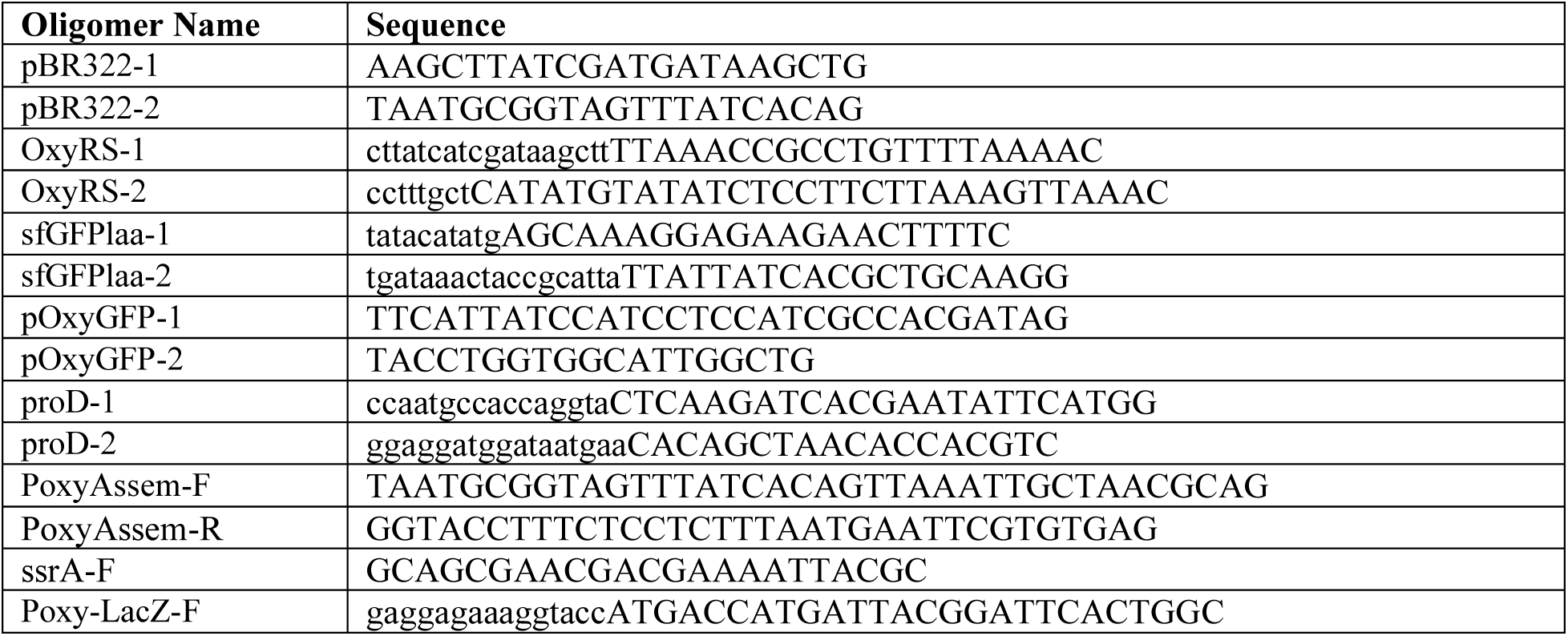
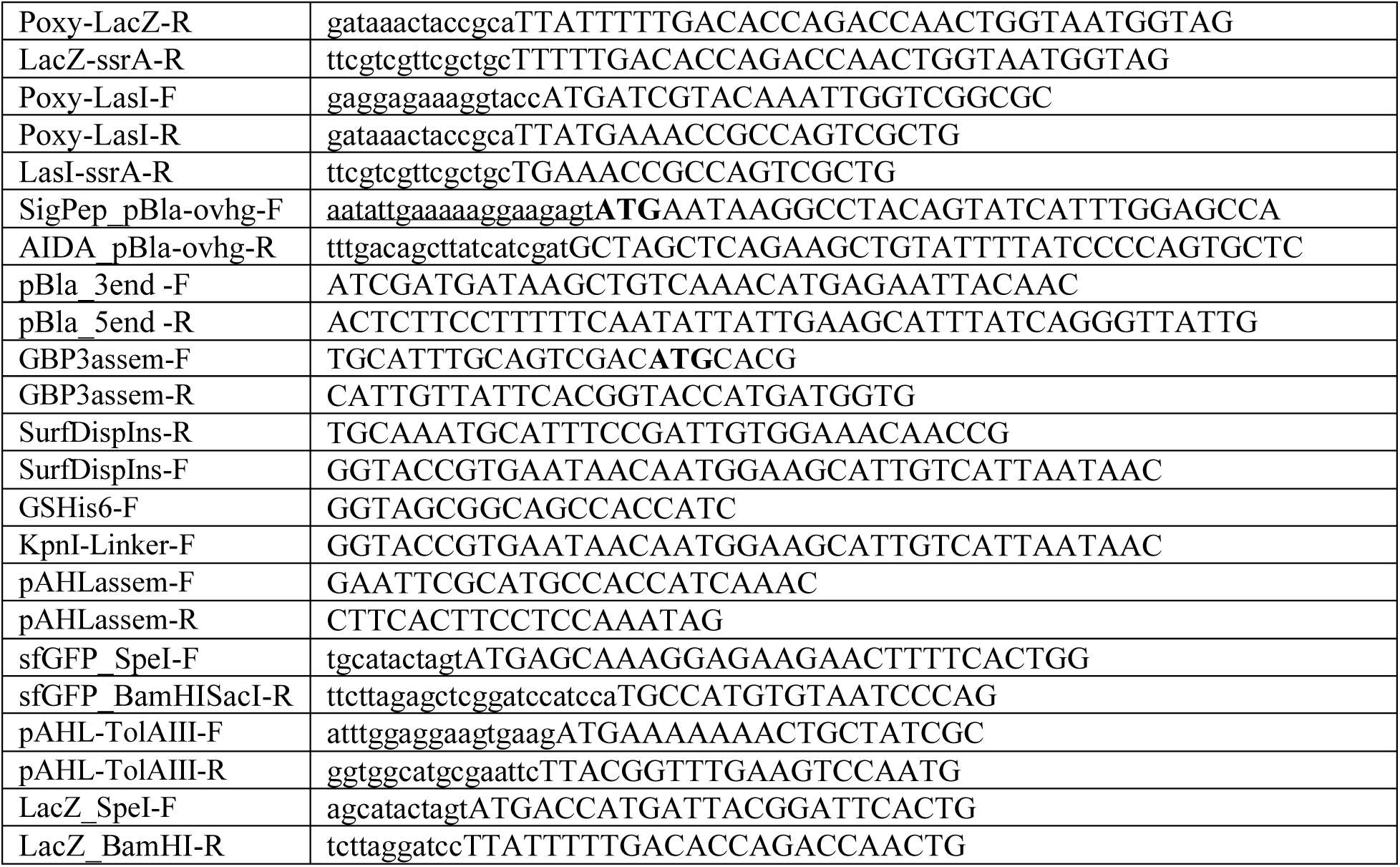
Primers used in this study.

**Supplementary Table 5.** Rate constants for modeling protein kinetics.

| Rate Constant | Value |
| --- | --- |
| $k_1$ | 2.2 OD<sup>-1</sup> · min<sup>-1</sup> |
| $k_2$ | 0.015 μM<sup>-1</sup> · min<sup>-1</sup> |
| $k_3$ | 14 min<sup>-1</sup> |
| $k_4$ | 0.018 [OxyR(o)]<sup>-1</sup>min<sup>-1</sup> |
| $k_{5\_gfp}$ | 0 |
| $k_{5\_βgal}$ | 0.00046 min<sup>-1</sup> |
| $k_{5\_lasI}$ | 0.0007 min<sup>-1</sup> |
| $k_{5\_AHL}$ | 2.01 min<sup>-1</sup> |
| $k_{6\_gfp}$ | 0.007 min<sup>-1</sup> |
| $k_{6\_βgal}$ | $k_{double}$ |
| $k_{6\_lasI}$ | 0.007 min<sup>-1</sup> |
| $k_{S\_gfp}$ | 0.4 |
| $k_{S\_βgal}$ | 0.3 |
| $k_{S\_lasI}$ | 0.3 |
| $k_{double}$ | 0.0115 min<sup>-1</sup> |
| $k_{7\_gfp}$ | 49,082 |
| $k_{7\_lacZ}$ | 73,170.73 |
| $k_{7\_lasI}$ | 22,000 |
| $K_{7\_AHL}$ | 67.63 |
| $\tau$ | 2.47 |
| $\sigma$ | 0.35 |
| $k_{8\_GFP}$ | 0.1 |

**Supplementary Table 6.**
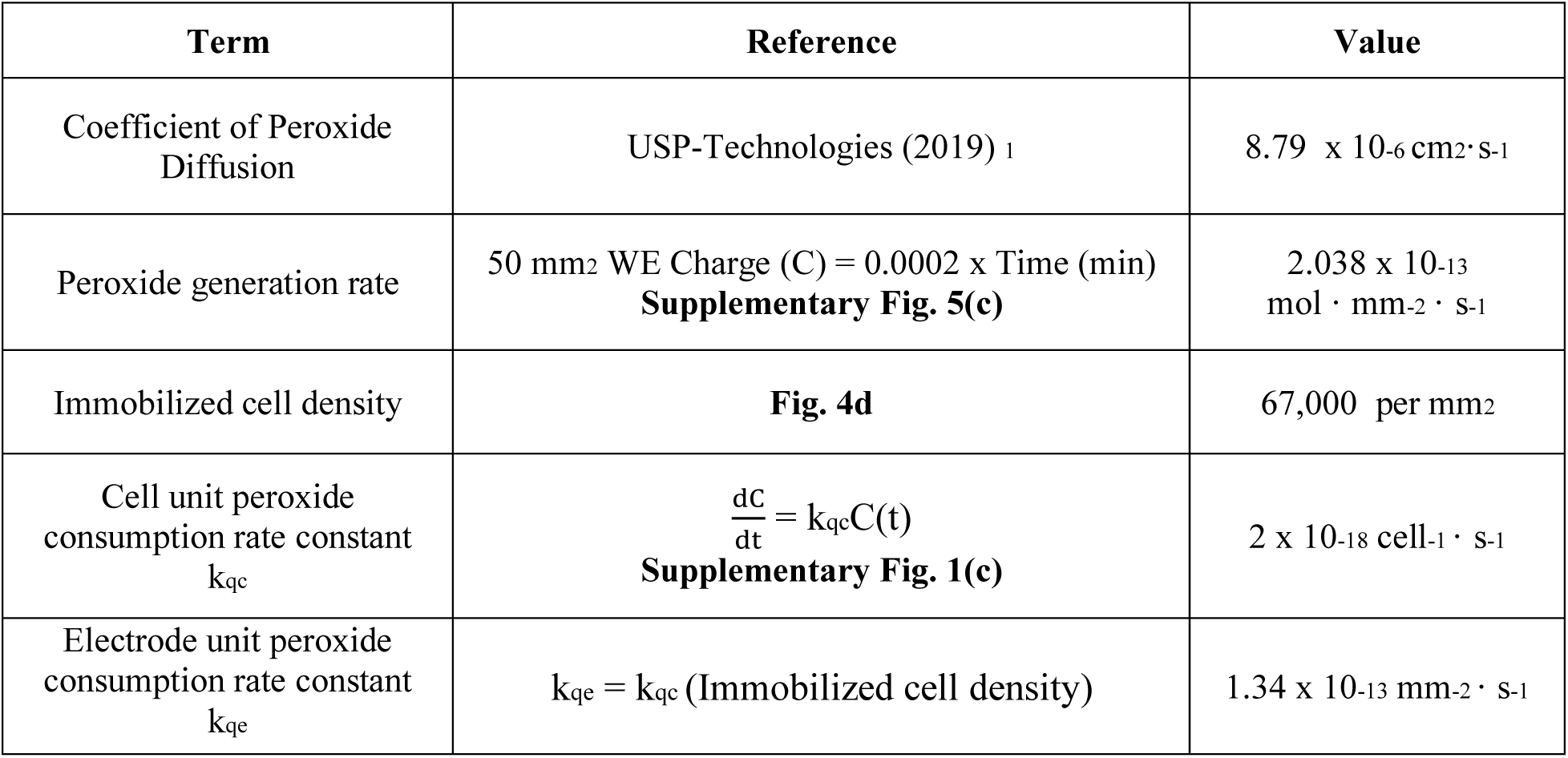
Parameters for modeling peroxide flux.

| Term | Reference | Value |
| --- | --- | --- |
| Coefficient of Peroxide Diffusion | USP-Technologies (2019) <sup>1</sup> | $8.79 \times 10^{-6} \text{ cm}^2 \cdot \text{s}^{-1}$ |
| Peroxide generation rate | 50 mm<sup>2</sup> WE Charge (C) = 0.0002 x Time (min)<br>**Supplementary Fig. 5(c)** | $2.038 \times 10^{-13} \text{ mol} \cdot \text{mm}^{-2} \cdot \text{s}^{-1}$ |
| Immobilized cell density | **Fig. 4d** | 67,000 per mm<sup>2</sup> |
| Cell unit peroxide consumption rate constant<br>$k_{qc}$ | $\frac{dC}{dt} = k_{qc}C(t)$<br>**Supplementary Fig. 1(c)** | $2 \times 10^{-18} \text{ cell}^{-1} \cdot \text{s}^{-1}$ |
| Electrode unit peroxide consumption rate constant<br>$k_{qe}$ | $k_{qe} = k_{qc} (\text{Immobilized cell density})$ | $1.34 \times 10^{-13} \text{ mm}^{-2} \cdot \text{s}^{-1}$ |

